# Characterizing the nuclear and cytoplasmic transcriptomes in developing and mature human cortex uncovers new insight into psychiatric disease gene regulation

**DOI:** 10.1101/567966

**Authors:** Amanda J. Price, Taeyoung Hwang, Ran Tao, Emily E. Burke, Anandita Rajpurohi, Joo Heon Shin, Thomas M. Hyde, Joel E. Kleinman, Andrew E. Jaffe, Daniel R. Weinberger

**Author notes:** ***Corresponding Author:*** Daniel R. Weinberger, 855 N Wolfe St, Ste 300; Baltimore MD 21205.

## Abstract

Transcriptome compartmentalization by the nuclear membrane provides both stochastic and functional buffering of transcript activity in the cytoplasm and has recently been implicated in neurodegenerative disease processes. Although many mechanisms regulating transcript compartmentalization are also prevalent in brain development, the extent to which subcellular localization differs as the brain matures has yet to be addressed. To characterize the nuclear and cytoplasmic transcriptomes during brain development, we sequenced both RNA fractions from homogenate prenatal and adult human postmortem cortex using PolyA+ and RiboZero library preparation methods. We find that while many genes are differentially expressed by fraction and developmental expression changes are similarly detectable in nuclear and cytoplasmic RNA, the compartmented transcriptomes become more distinct as the brain matures, perhaps reflecting increased utilization of nuclear retention as a regulatory strategy in adult brain. We examined potential mechanisms of this developmental divergence including alternative splicing, RNA editing, nuclear pore composition, RNA binding protein motif enrichment, and RNA secondary structure. Intron retention is associated with greater nuclear abundance in a subset of transcripts, as is enrichment for several splicing factor binding motifs. Finally, we examined disease association with fraction-regulated gene sets and found nuclear-enriched genes were also preferentially enriched in gene sets associated with neurodevelopmental psychiatric diseases. These results suggest that although gene-level expression is globally comparable between fractions, nuclear retention of transcripts may play an underappreciated role in developmental regulation of gene expression in brain, particularly in genes whose dysregulation is related to neuropsychiatric disorders.

## Introduction

Human brain development is marked by precisely-timed changes in gene expression across the lifespan, with the most dramatic changes occurring at the prenatal to postnatal transition (1–3). One factor governing these changes is the compartmentalization of the transcriptome by the nuclear membrane into nuclear and cytoplasmic fractions. A snapshot of each RNA compartment composition captures factors of both chance and purpose: since most splicing of pre-mRNA occurs co-transcriptionally in the nucleus (4, 5), for instance, pre-mRNA and longer genes that take more time to be transcribed and exported are often overrepresented in the nucleus compared to cytoplasm (6–8). Recent studies have also highlighted the role of the nuclear membrane as a transcriptional noise buffer, filtering stochastic bursts of gene expression from the cytoplasm by retaining selected mature mRNA transcripts in the nucleus (9, 10). Still, nuclear retention can also be used by the cell to regulate the timing of cytoplasmic activity of a transcript (11, 12) as well as perform quality control by sequestering aberrant transcripts in the nucleus and targeting them for degradation.

Interestingly, many of these RNA trafficking mechanisms are particularly prevalent in brain cells and some have been found to play a role in brain development. For example, alternative splicing—particularly intron retention—has been shown to regulate RNA localization as a means to suppress lowly- and aberrantly-expressed transcripts (13, 14). Intron retention is common in neuronal lineages and serves to down-regulate genes involved in other lineage fates during neuronal differentiation (14–16). RNA editing is also developmentally regulated in human brain, with a subset of editing sites associated with neuronal maturation (17). In at least one example, RNA editing was also shown to regulate activity-dependent nuclear transcript retention, although global characterization of RNA editing patterns by subcellular fraction shows that RNA editing is not broadly necessary for nuclear retention (11, 18). Likewise, nuclear pore composition is also developmentally dynamic, and the RNA binding proteins (RBPs) that regulate many of the above processes and shepherd RNA from the nucleus to the cytoplasm are also under developmental control (19, 20).

Although nuclear and cytoplasmic transcriptomes have been assessed using ***in vitro*** models, subcellular fractions have not yet been characterized in human cortical brain tissue and not across age dimensions. Given the increasingly frequent use of nuclear RNA in single cell and cell population-based studies of human brain as a result of the difficulty of dissociating frozen post-mortem brain tissue to a single cell suspension (21–23), understanding the compositional differences between RNA compartments over human brain development would also help inform future studies using nuclear RNA without a comparable cytoplasmic fraction. Given also that disruption of proper nucleocytoplasmic transport of proteins and RNA is increasingly implicated as a mechanism in normal aging as well as neurodegenerative disorders such as fronto-temporal dementia, Alzheimer’s disease, Huntington’s disease, and amyotrophic lateral sclerosis (24–27), characterizing the dynamics of RNA localization between the fractions may help clarify what role if any nuclear sequestration of transcripts may play in the etiology of brain disorders with developmental components.

To address these questions in human cortical tissue, we characterized the nuclear and cytoplasmic transcriptomes in early developing and mature prefrontal cortex using two RNA sequencing library preparation methods and the same homogenate tissue dissections for each—matching similarly aged samples for tissue-level cellular composition—and examined distributions of gene sets associated with neurodevelopmental, neurodegenerative, and psychiatric disorders. We show that although many genes are differentially expressed by subcellular fraction, developmental differences in gene expression are similarly detectable in nuclear and cytoplasmic RNA. Despite this overall similarity, RNA compartments become more distinct as the brain matures, perhaps as a result of greater use of nuclear retention as a regulatory strategy. We also explored potential mechanisms of gene expression regulation by fraction and assessed the relationship between nuclear pore gene expression, intron retention, RNA editing, RBP motif enrichment, and RNA secondary structure and RNA subcellular localization patterns, with splicing decisions potentially emerging as the most salient contributor behind differences in fraction expression patterns. Finally, we found nuclear-enriched genes in both prenatal and adult cortex to be enriched for psychiatric disorder gene sets, and identify intron retention patterns and splicing-associated RBPs that may contribute to their nuclear association. These results describe the fractionated transcriptome in great detail and point to a potentially underappreciated role for nuclear compartmentalization in psychiatric disease gene regulation.

## Results

We sequenced nuclear and cytoplasmic RNA isolated from three prenatal and three adult human brains, constructing libraries using two distinct strategies for depleting ribosomal RNA, the dominant RNA species in total RNA (PolyA+ and Ribo-Zero; Fig. S1, Table S1). Together, PolyA+ library preparation (selecting polyadenylated transcripts via a pull-down step) and Ribo-Zero library preparation (relying on an rRNA depletion step) capture the transcriptomic diversity in these subcellular compartments in developing human brain due to their respective preferences for mature mRNA and non-polyadenylated transcripts (e.g., ncRNA or pre-mRNA, Fig. S2A)(28–30). One adult nuclear Ribo-Zero sample failed quality control and was discarded and two prenatal cytoplasmic PolyA+ samples that had higher read depth were down-sampled to a comparable depth. In total, we profiled 43,610 Ensembl genes that were expressed across any of these eight groups of samples (Adult/Cytoplasm, Adult/Nucleus, Prenatal/Cytoplasm, and Prenatal/Nucleus with both library types; see Fig. S1). The quality of fractionation was confirmed by determining that two genes known to localize either to the nucleus (e.g., *MALAT1*) or cytoplasm (e.g., *ACTB*) (9) were significantly enriched in the appropriate compartment (p-value adjusted by False Discovery Rate (FDR)<0.01; Fig. S2B), although prenatal samples showed less enrichment for these genes than adult samples (FDR=1.2e-6 and FDR=9.6e-9 for *ACTB* and *MALAT1* in adult, versus FDR=0.17 and FDR=0.44 in prenatal, respectively). A similar amount of RNA was collected from the nuclear compared to cytoplasmic fraction in prenatal and adult samples, also suggesting adequate fractionation (FDR>0.05; Fig. S2C).

### Developmental gene expression changes in human cortex are similarly detectable in nuclear and cytoplasmic RNA

We first defined the RNA content differences between subcellular fractions and replicated many characteristics that have previously been described (4–7, 9, 31, 32). Genes that are significantly more abundant in the nucleus were overall longer than genes more abundant in the cytoplasm, perhaps due to the longer temporal requirement for transcription and diffusion through the nuclear pore (Fig. S2D). The proportion of reads aligning to introns was greater in the nucleus than the cytoplasm in both PolyA+ and Ribo-Zero samples (t>4.7, FDR<5.96e-3), indicating a greater proportion of immature pre-mRNA or unannotated transcripts (Fig. S2E). Fraction-regulated genes were expressed in a variety of cell types: over a quarter (25.9-29.3%) of both cytoplasmic and nuclear-enriched genes were highest expressed in neurons from prior single-cell RNA-seq data (33), while 18.1-19.7% were highest expressed in astrocytes, 11.1-12.7% were highest expressed in oligodendrocytes or their precursors, 4.0-4.5% were highest in microglia, and 4.6-6.1% were highest in epithelial cells (Fig. S2F). While the majority (83.5%) of all fraction-regulated genes were protein-coding (Fig. S2G), a larger proportion of genes enriched in the nuclear fraction were non-coding than those enriched in the cytoplasm (OR=0.25, p=2.2e-16). Because Ribo-Zero libraries do not require polyadenylation for sequencing, a greater proportion of differentially expressed genes across fractions were non-coding in Ribo-Zero samples (Fig. S3A-B).

Expression patterns were overall similar between fractions at the gene level. Principal component analysis showed that sample age and library type were the largest contributors to transcriptomic diversity, explaining 53% and 35% of the variance, respectively (Fig. 1A). Developmental expression trajectories were highly correlated between the fractions (*ρ*=0.89, t = 335.8, p<2.2e-16; Fig. 1B), and 41-63% of genes differentially expressed across development as measured in the four fraction/library groups were detected in all groups (Fig. 1C). In summary, developmental changes in gene expression measured in the nuclear compartment were comparable to the changes detected in the cytoplasmic fraction, supporting the conclusion that nuclear gene expression is representative of gene expression at the cellular level.

**Figure 1:**
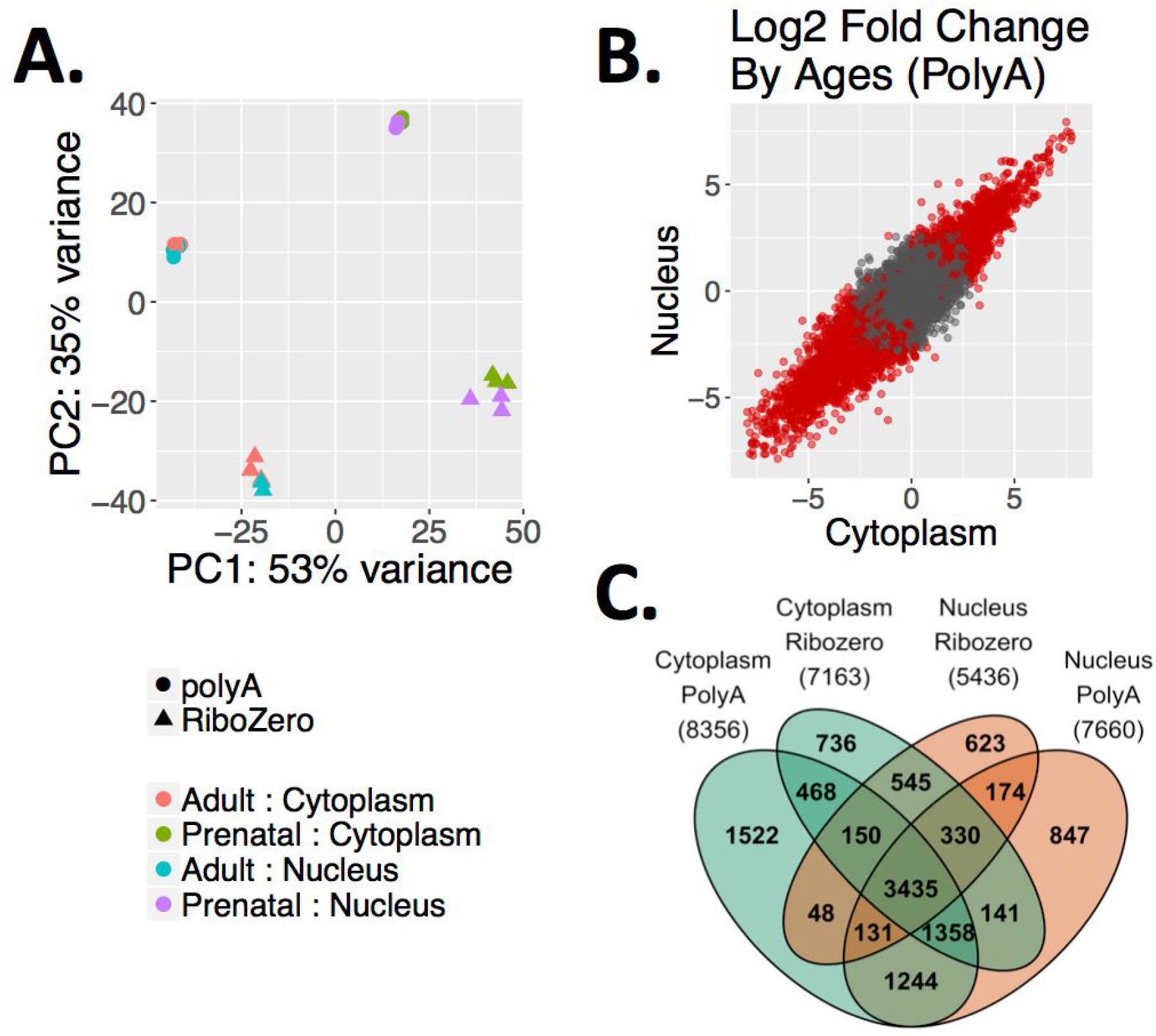
Developmental gene expression changes in human cortex are similarly detectable in nuclear and cytoplasmic RNA. **(A)** Principal component analysis. PC1 separates the samples by age and PC2 separates the samples by library type so that nuclear and cytoplasmic samples from the same donors cluster together. **(B)** log_2_ fold change of gene expression across age measured in cytoplasmic and nuclear RNA (PolyA+ library preparation shown only). Red dots indicate significantly differentially expressed genes (FDR≤0.05). **(C)** Venn diagram of differentially expressed genes by age (FDR≤0.05; abs(log2 fold change)≥1) measured in both RNA fractions and library types. The total number of genes for each group is listed in parentheses.

### Prenatal and adult human cortex show distinct patterns of RNA localization across the nuclear membrane

We next examined the relationship between developmental stage and gene expression by fraction and found that prenatal and adult cortex exhibited substantially different RNA localization patterns across the nuclear membrane. We identified 1,894 genes differentially expressed by fraction in adult cortex, but only 40 genes differentially expressed in prenatal cortex in the PolyA+ samples (Fig. 2A, abs(Log_2_ Fold Change)≥1; FDR≤0.05). This localization pattern difference was also seen in Ribo-Zero samples (Fig. S3C-D). Interestingly, unlike in adult samples where differentially expressed genes by fraction (FDR≤0.05) were equally likely to be nuclear or cytoplasmic, most fraction-regulated genes in prenatal samples were more abundant in the nuclear compartment (Table S2). Despite fewer genes being differentially expressed by fraction in prenatal cortex, subcellular expression patterns were correlated between prenatal and adult (*ρ*=0.60, t=125.9, p<2.2e-16; Fig. 2B).

**Figure 2:**
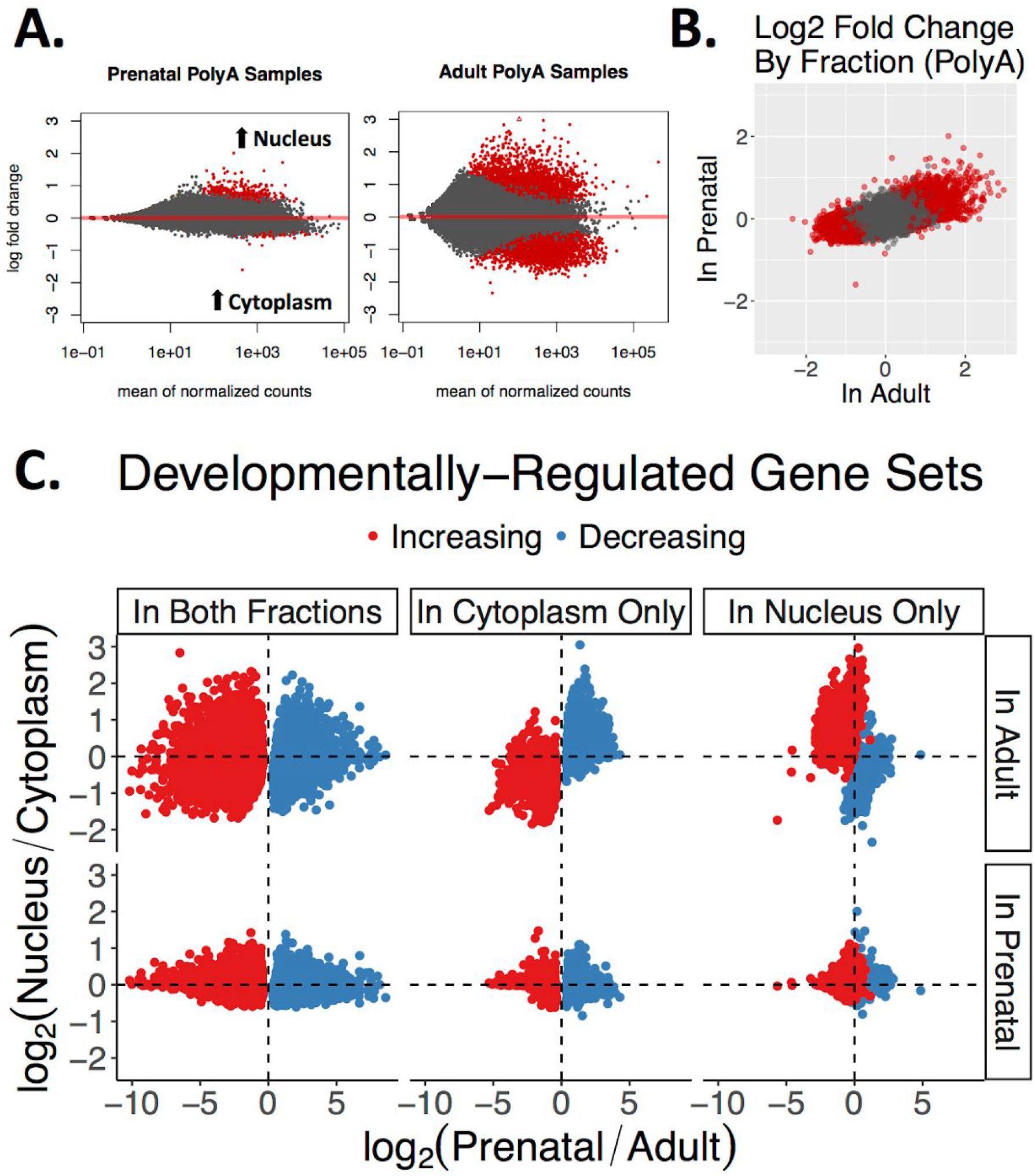
Prenatal and adult human cortices show distinct patterns of RNA localization across the nuclear membrane. **(A)** MA plots of prenatal and adult gene expression differences across fraction. Red dots indicate FDR≤0.05. **(B)** Log_2_ fold change of gene expression across fraction in adult PolyA+ samples plotted against prenatal PolyA+ samples. Red dots indicate significantly differentially expressed genes (FDR≤0.05). **(C)** Log_2_ fold change between fraction and age for six developmentally-regulated gene sets: colors indicate developmental expression trajectory direction, and columns indicate the fraction in which the developmental expression difference is detected. The y-axis shows the log_2_ fold change between fractions stratified by the age in which the fraction differences are measured, and the x-axis shows the log_2_ fold change between ages as measured in cytoplasm. While these developmentally-regulated gene sets are similar across prenatal fractions, down-regulated genes in adult are more sequestered in the nuclear compartment.

The association of fraction expression with changing developmental expression depended on the fraction in which the developmental changes were measured. This can be seen when exploring the differences in fraction expression in six developmentally regulated gene sets (Fig. 2C; Table S3). Genes with changing developmental trajectories detected in both fractions showed no difference in expression between fractions. In contrast, genes with decreasing expression from prenatal to adult when measured in cytoplasmic RNA (i.e., developmentally decreasing expression in cytoplasm only) were more likely to be higher expressed in adult nucleus than adult cytoplasm (OR=38.0, FDR=1.9e-20), while developmentally decreasing genes detected in nuclear RNA were less likely to be expressed in adult nucleus compared with adult cytoplasm (OR=0.071, FDR=1.7e-10). In other words, prenatally enriched RNA may be relatively sequestered in the nuclear compartment in adult cortex to regulate the translation of these more fetal-relevant genes in adulthood. Likewise, genes with greater expression in adult than prenatal cytoplasm were less expressed in adult nucleus (OR=0.038, FDR=4.7e-24), while developmentally increasing genes in nuclear RNA were more likely to be greater expressed in adult nucleus than cytoplasm (OR=19.3, FDR=5.5e-10). This pattern was not seen in prenatal RNA fractions, as expression differences between fractions were more muted in this age. Taken together, these patterns suggest an inverted relationship between developmental gene expression changes and subcellular compartmentalization, where genes with decreasing expression across development are progressively more likely to be retained in the adult nucleus.

### A subset of introns regulated by fraction and age is associated with decreasing developmental expression and increasing nuclear localization

We next sought to provide context for these RNA localization patterns by examining several mechanisms associated with RNA transport across the nuclear membrane. Because alternative splicing—particularly intron retention—has been implicated as a mechanism of localization of transcripts within the cell (13, 14) and can play a role in regulating developmental gene expression (15, 16), we next characterized alternative splicing across the PolyA+ samples. The proportion of reads spanning splice junctions was not significantly different between the nuclear and cytoplasmic fractions in the PolyA+ samples, as pre-mRNAs were overall depleted by PolyA+ selection (t=-1.0, FDR=0.34; Fig. 3A), suggesting that much of the RNA captured via PolyA+ selection is already processed. In contrast, the proportion of spliced alignments was much lower in nuclear than cytoplasmic Ribo-Zero samples, suggesting greater representation of pre-mRNA in the libraries (t=-4.3, FDR=0.016), and an overall relative reduction of mature RNA in the nucleus when depleting rRNA. All following splicing analyses were therefore performed in the PolyA+ samples.

**Figure 3:**
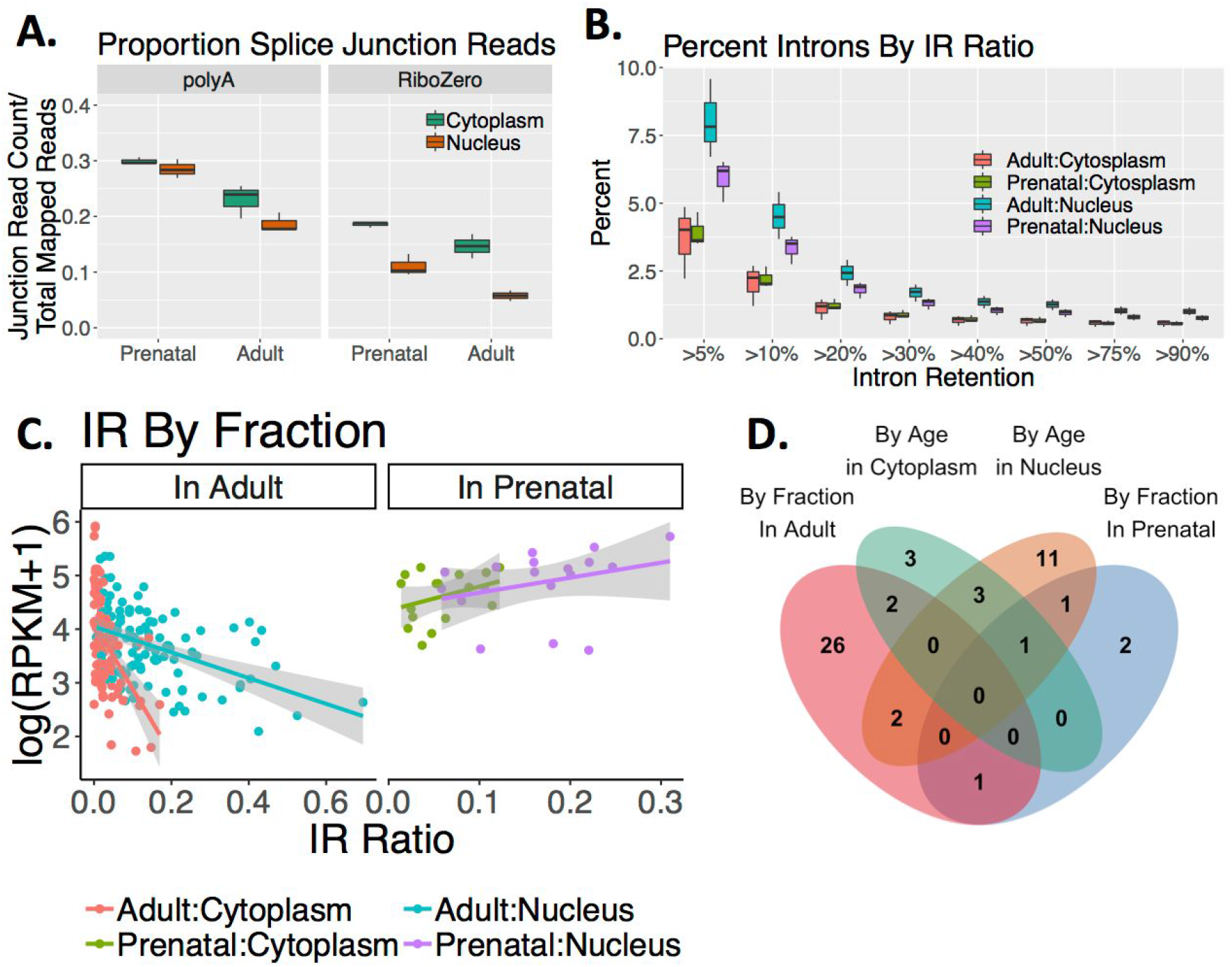
Intron retention patterns in prenatal and adult human cortex associate with RNA distribution. **(A)** Proportion of junction reads per sample by group. Prenatal cortex has a higher proportion of splice junctions than adult, and although RiboZero samples have a lower proportion of junctions in nuclear samples, PolyA+ samples show non-significantly different proportions between fractions. **(B)** Percentage of introns passing QC in all groups with an intron retention (IR) percentage above each threshold in at least one sample of each group. Introns in nuclear samples had greater IR ratios than cytoplasmic samples (t=69.5, FDR=0). **(C)** The IR ratio of introns differentially retained by fraction when measured in adult and prenatal samples, plotted by gene expression measured as the log of the reads per kilobase per million mapped (RPKM) plus one. Samples are colored by their age:fraction group. **(D)** Overlap of genes containing introns that were differentially retained by fraction or age (FDR≤0.05).

After surveying the diversity of splicing events in both RNA fractions and ages (Fig. S4A-B), we focused on intron retention, examining 166,661-173,125 introns per sample after filtering that represented 15,345-15,389 unique genes (Fig. 3B). Greater intron retention was neither associated with the localization of a transcript by fraction overall (Fig. S4C), nor correlated with developmental changes to gene expression overall (Fig. S4D). However, a small subset of individual introns were differentially retained in a transcript by fraction and age and showed distinct relationships with gene expression that suggest differing activities by fraction and age (Fig. S4E). For instance, fraction-regulated introns in adult but not prenatal samples were negatively correlated with the expression of their host genes (ρ<-0.44, t<-5.0, FDR<1.8e-05; Fig. 3C). One interpretation of this result is that increasing intron retention downregulates expression in adult but not prenatal by signaling for nuclear degradation in adult of these genes. Indeed, increasing IR is positively correlated with nuclear enrichment of the intron-containing genes (ρ>0.45, t>3.9, FDR<1.0e-03; Fig. S4F). Differentially retained introns by age showed no relationship between IR and expression levels except in prenatal cytoplasm, where increasing IR was also correlated with decreasing expression (ρ=-0.69, t=-5.1, FDR=1.2e-04; Fig. S4G).

Fraction and age-associated introns often were included in the same genes, emphasizing the connection between development and RNA localization. Genes including significantly differentially retained introns by fraction (FDR≤0.05) were more likely also to include significantly differentially retained introns by age (OR=88.4, FDR=6.5e-04). Developmentally-regulated introns were also more likely to be in genes that were significantly differentially expressed by fraction and vice versa (OR>2.8, FDR<0.032; Fig 3D). Interestingly, these intron-containing genes—particularly those containing nuclear-enriched introns in adult and prenatal-enriched introns in nuclear RNA—were enriched for ontology terms associated with neuron-specific cellular compartments (FDR≤0.05; Fig. S5A-B). While most introns are not associated with gene expression patterns by fraction or age, a subset showed a decreasing expression of their host genes, often genes implicated in both processes as IR increases.

### Highly edited gene products contain RNA editing sites unique to an age/fraction group that associate with expression levels

We next profiled RNA editing across subcellular fractions in prenatal and adult cortex. We identified 3,064-5,840 editing sites per sample, finding 25,051 unique sites across the dataset (Fig S6A-E). Of these, 75.5% were A-to-I edited sites, the most common editing pattern (Appears as A:G or T:C in our sequencing data; Fig. 4A). Of the 18,907 A-to-I edited sites, only 1,025 were shared by all four groups, representing 729 unique genes (Fig. 4B). To assess the relationships between subcellular localization and RNA editing, we first assessed editing rate changes across fraction and age in the 1,025 A-to-I sites shared among the four groups. The distribution of unadjusted p-values suggested that age but not fraction influenced overall editing rates (Fig. S6F).

**Figure 4:**
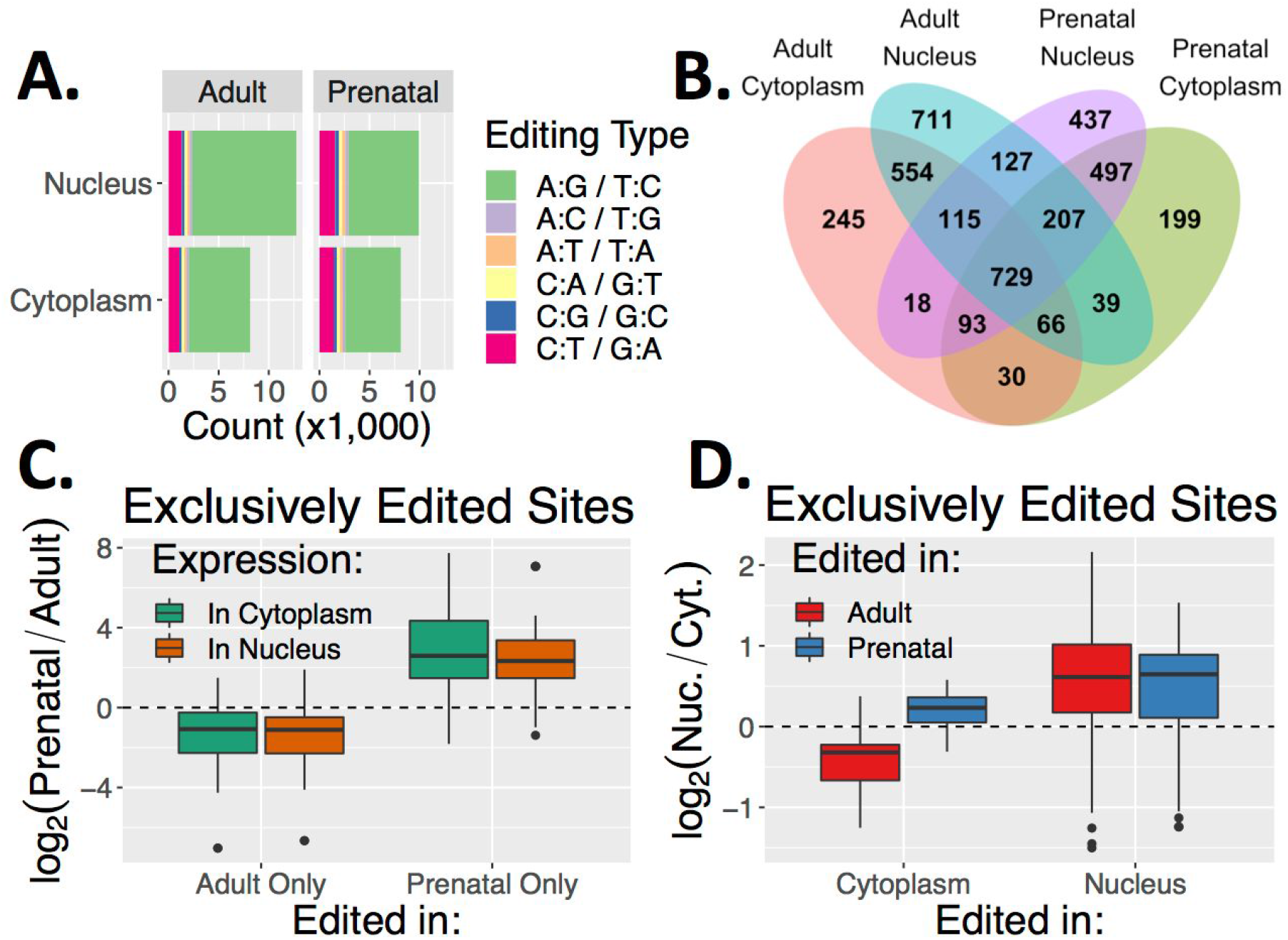
RNA editing sites unique to an age/fraction group associate with expression levels. **(A)** Number of unique editing sites identified in each group stratified by editing context. **(B)** Venn diagram demonstrating the overlap of edited genes between groups of fraction and age samples. **(C)** Log_2_ fold change of expression by age in genes that include an editing site present in all adult but no prenatal samples (Adult Only) or all prenatal but no adult samples (Prenatal Only), as measured in cytoplasmic and nuclear RNA. **(D)** Log_2_ fold change of expression by fraction as measured in adult samples, in genes that include an editing site present in all cytoplasmic but no nuclear samples (right) or all nuclear and no cytoplasmic samples (left) in either adult (red) or prenatal (blue) samples.

Because these 1,025 sites reflected only 5.4% of the A-to-I editing sites, we focused on the sites found consistently in every sample of one of the four PolyA+ groups from above and never in a contrasting group, thinking that these sites may be functionally important to that group’s compartment and age time point (Table S4; summarized in Table 1). Genes containing an editing site in this subset had significantly more edited sites than other genes (t>3.3, FDR<6.9e-03). These edited genes were also higher expressed in the compartment or age containing the edited site: for instance, genes that were significantly greater expressed in adult samples were enriched for sites consistently and exclusively edited in adult samples (OR=8.9, FDR=6.3e-19), while genes that were significantly greater expressed in prenatal cortex were enriched for sites consistently and exclusively edited in prenatal samples (OR=25.9, FDR=2.1e-26, Fig. 4C). The editing sites that were found in all nuclear but no cytoplasmic samples (ie., the 159 in adult and 65 in prenatal nuclear RNA from Table 1) were more likely to occur in genes that were significantly higher expressed in nuclear RNA than other editing sites (OR>2.9, FDR<2.3e-02; Fig. 4D; Fig. S6G), raising the possibility that these nuclear-unique editing sites help in signaling nuclear sequestration. Although these editing site subsets were found exclusively in one compared to a contrasting group, 86.49-100% of the introns and 96.55-100% of the exons including these editing site were expressed unedited in the contrasting groups, meaning the sequence being edited was usually expressed unedited in other fractions and ages.

**Table 1:**
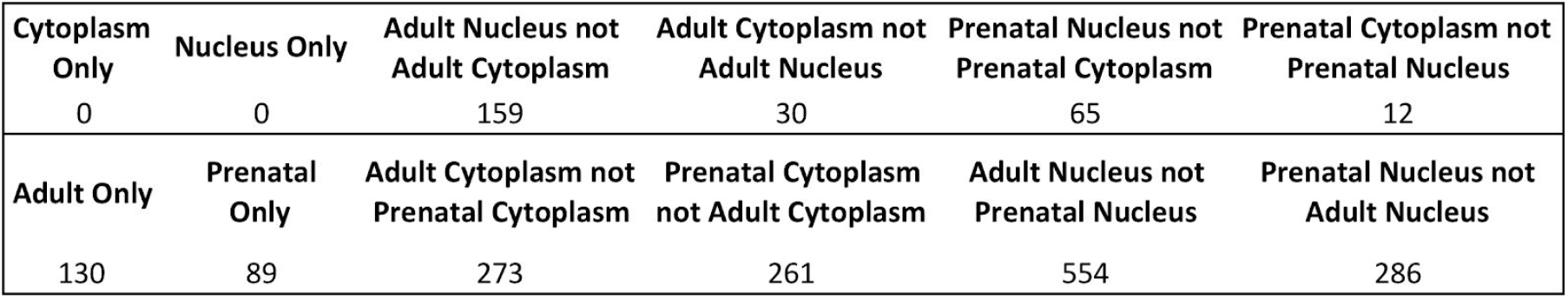
Unique RNA Editing Sites By Fraction and Age. The number of RNA editing sites found in all samples of a group but not found in a contrasting group. For age and fraction individually, the site must be present in all six PolyA+ samples, while subsets of fraction and age must be present in the three samples. For example, 159 editing sites were found in all three adult nuclear RNA samples but were not detected in any adult cytoplasmic RNA samples.

|  |  |  |  |  |  |
| --- | --- | --- | --- | --- | --- |
| **Cytoplasm Only** | **Nucleus Only** | **Adult Nucleus not Adult Cytoplasm** | **Adult Cytoplasm not Adult Nucleus** | **Prenatal Nucleus not Prenatal Cytoplasm** | **Prenatal Cytoplasm not Prenatal Nucleus** |
| 0 | 0 | 159 | 30 | 65 | 12 |
| **Adult Only** | **Prenatal Only** | **Adult Cytoplasm not Prenatal Cytoplasm** | **Prenatal Cytoplasm not Adult Cytoplasm** | **Adult Nucleus not Prenatal Nucleus** | **Prenatal Nucleus not Adult Nucleus** |
| 130 | 89 | 273 | 261 | 554 | 286 |

### Nuclear-enriched genes contain motifs for distinct sets of RNA binding proteins and have more stably structured 3’UTRs

To examine the potential role of specific RBPs in RNA localization by fraction over brain development, we examined the enrichment of 1,157 sequence motifs corresponding to 140 RBPs in genes differentially regulated by fraction in adult and prenatal cortex (FDR≤0.05; Table S5). We found that of the 86-123 unique RBPs with motifs enriched in each of the four gene sets (e.g., cytoplasmic in adult, cytoplasmic in prenatal, nuclear in adult, and nuclear in prenatal; FDR≤0.05 for genes’ inclusion in the set, FDR≤0.01 for motif enrichment), the majority was shared across age and fraction (61.8-88.4% of RBPs per gene set; Fig. 5A). Many of the RBPs exclusively enriched in one fraction versus the other have previously been implicated in neurodevelopment, and the fraction in which they were enriched corresponded to the RBP’s most prevalent location of action. For instance, motifs for ELAVL2 and ELAVL3 are found exclusively in cytoplasmic-enriched genes compared to nuclear genes, and both RBPs act in neuronal dendrites to regulate neuronal development and function (34). Likewise, AGO1 and AGO2 function in the cytoplasm as part of the RISC complex, and are both are exclusively enriched in cytoplasmic genes. On the other hand, motifs for many nuclear-acting factors including CELF4 and PTBP2—RBPs that are involved in splicing, mRNA stability and transport and are implicated in neuronal differentiation, plasticity and transmission—are exclusively enriched in nuclear-enriched genes (34). In fact, many of the nuclear-enriched RBPs were especially relevant to splicing decisions occurring in neuronal nuclei, such as the neuron-specific splicing factors RBFOX1 and RBFOX2. RBFOX1 particularly is implicated in developmental control of neuronal excitability and synaptic transmission, and has been shown to regulate the splicing patterns of other RBPs, such as hnRNPH1, ELAVL2, and PTBP (34). Given the localization and functional implications of the enriched RBPs, it is possible that RBP binding plays a part in the differential expression of these genes across subcellular fractions.

**Figure 5:**
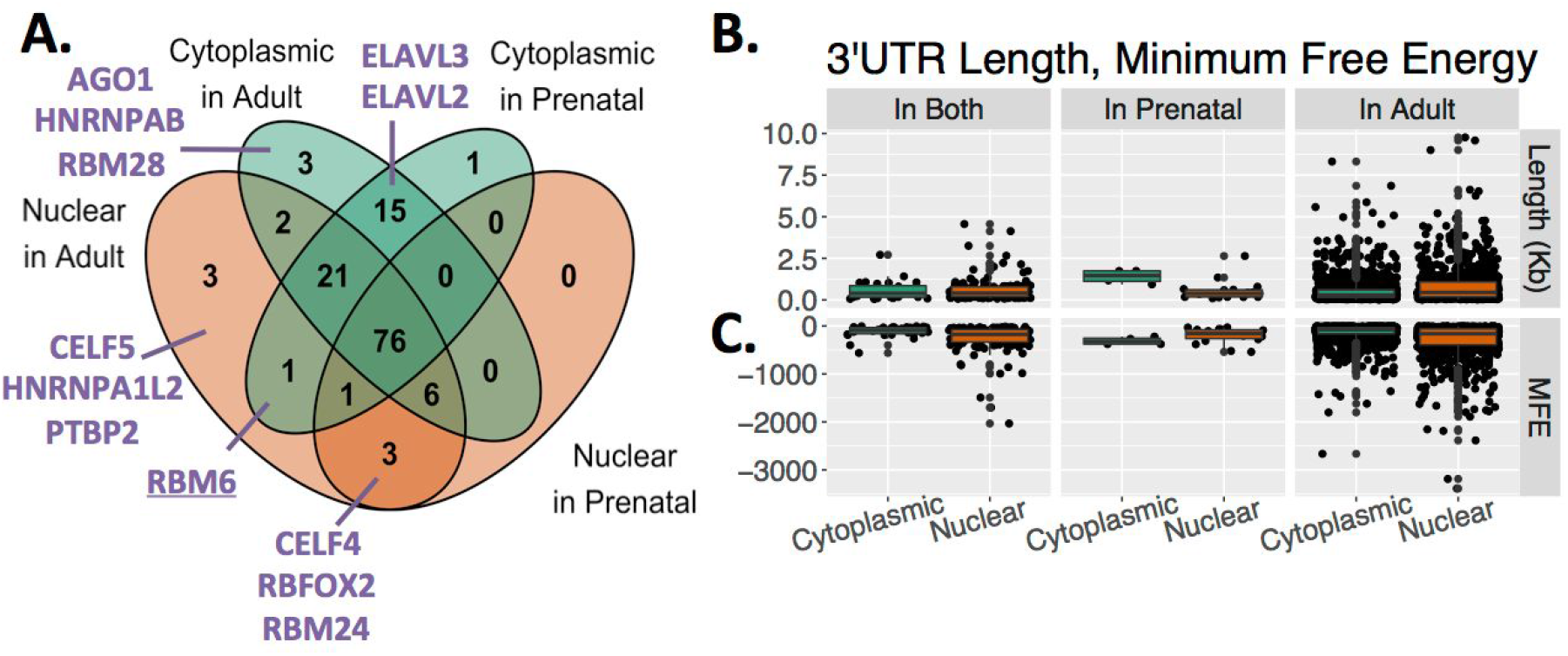
RNA binding protein motif enrichment, 3’UTR length and predicted secondary 3’UTR structure of fraction-regulated genes. **(A)** Overlap of RBPs with sequence motifs found (FDR≤0.01) in four groups of genes differentially expressed by fraction measured in adult and prenatal samples (FDR≤0.05). RBPs discussed in the text are listed in purple. **(B)** Length in kilobases of the highest expressed 3’UTR for each gene differentially expressed by fraction in both adult and prenatal, only prenatal, or only adult brain. **(C)** Minimum predicted free energy of the secondary structure for the above 3’UTRs.

Sequence recognition motifs are but one element governing RBP binding, and recent work has shown that other factors such as RNA secondary structure are more consequential in determining putative binding patterns, especially conjointly with recognition motifs (35). 3’UTR sequence is particularly critical to acting as an anchor for these RBPs (36). Because of this, we measured the minimum free energy (MFE) of the predicted secondary structure for the highest expressed 3’UTR for each gene differentially expressed by fraction and found that genes higher expressed in nuclear RNA in both ages or in adult only had significantly lower MFE than cytoplasmic genes, meaning that nuclear-enriched gene 3’UTRs were predicted to be more thermodynamically stable than 3’UTRs of cytoplasmically-enriched genes (t<-3.9, FDR<2.7e-04; Fig. 5B). This result was not necessarily a factor of differing 3’UTR length, as 3’UTRs of nuclear genes from both ages were of comparable size (t=0.36, FDR=0.72), while adult-specific nuclear-enriched genes had longer 3’UTRs (t=7.3, FDR=2.7e-12) and prenatal-specific nuclear enriched genes had shorter 3’UTRs (t=-3.4, FDR=1.8e-02; Fig. 5C). Having a more stable 3’UTR RNA secondary structure in the 3’UTR may therefore potentially contribute to the subcellular RNA localization patterns of these nuclear-enriched genes.

### A subset of increasingly expressed nuclear pore genes is associated with the Nup107-160 nuclear pore subcomplex

An integral facet of nuclear transport are the nuclear pore complexes (NPCs) that facilitate passage through the membrane. Although NPC structures are extremely long-lasting, often persisting for the lifespan of a cell (37), many components of the NPCs can be exchanged according to the requirements of the cell, including during neuronal development (19). We therefore measured developmental expression trajectories of genes associated with NPCs and identified 42 NPC genes with decreasing and nine NPC genes with increasing expression in either subcellular RNA compartment as the brain matures (FDR≤0.05; see the final column of Table S3).

Several of the NPC factors with increasing expression are involved with the Nup107-160 nuclear pore subcomplex, a conserved component of the NPC that is involved in mRNA export and interacts with chromatin to establish new NPCs after mitosis (38). For instance, *SEH1L* (a.k.a. *Seh1*) is increasingly expressed in cortex and encodes a component of the Nup107-160 nuclear pore subcomplex (39). SENP2, an increasingly expressed SUMOylation protease, also interacts with the Nup107-160 nucleoporin subcomplex in addition to other NPC components (40). RanGAP1-RanBP2 is also increasingly expressed in cortex and is recruited by the Nup107-160 nuclear pore subcomplex to the kinetochores during mitosis (39). Although the association of the Nup107-160 nuclear pore subcomplex and RanGAP1 with proliferative phenotypes and mitosis (41) likely points to increasing expression due to increasing gliogenesis over the human lifespan, *RanGAP1* is higher expressed in human neurons than in glial subtypes in a publicly available single cell RNA-seq dataset, implicating neurons as well (FDR=2.1e-03; Fig. S7A)(33). *NUP107* and *NUP133*, main components of the Nup107-160 nuclear pore subcomplex, are not expressed in a cell type-specific way (Fig. S7B). Although NPC structures are stable in postmitotic neurons of adult cortex and developmental expression of these pore related genes did not explain other results, increasing expression of Nup107-160 subcomplex components and interactors may reflect additional roles of the subcomplex in these cells.

### Genes differentially expressed by fraction are overrepresented in gene sets associated with brain disease

We performed Disease Ontology (DO) Semantic and Enrichment analysis on the sets of genes differentially expressed by fraction and age, finding that genes with an interaction between fraction and age are enriched for genes associated with Alzheimer’s and Parkinson’s Diseases, in agreement with previous work implicating nucleocytoplasmic transport in neurodegenerative disease (Fig. S8A-B).

We then assessed the relationship between the groups of fraction-associated genes (see Table S2) with gene sets for neurodevelopmental, neurodegenerative, and psychiatric disorders curated from many sources, including genome-wide association (GWAS), copy number variation (CNV), and single nucleotide variation (SNV) studies (42) (Table S6). Neurodegenerative disease genes were enriched for cytoplasmic genes in adult cortex (OR=4.3, FDR=1.5e-3), while intellectual disability genes were enriched for cytoplasmic genes in both ages (OR=16.1, FDR=0.012), commensurate with their high expression in brain and the onset timing of each disorder. Interestingly, nuclear genes in both ages were enriched for genes associated with both Autism Spectrum Disorder (ASD; OR>4.9, FDR<4.0e-3; Fig S8C), as well as schizophrenia (SCZ; OR=6.5, FDR=0.014; Fig S8D). Bipolar Affective Disorder (BPAD) was also associated with genes greater expressed in nuclear RNA in adult cortex but not prenatal cortex (OR=3.1, FDR=1.5e-3; Fig 6A). Intellectual disability, syndromal neurodevelopmental disorder, and neurodegenerative disorder gene sets were neither enriched nor depleted for the nuclear-expressed genes.

**Figure 6:**
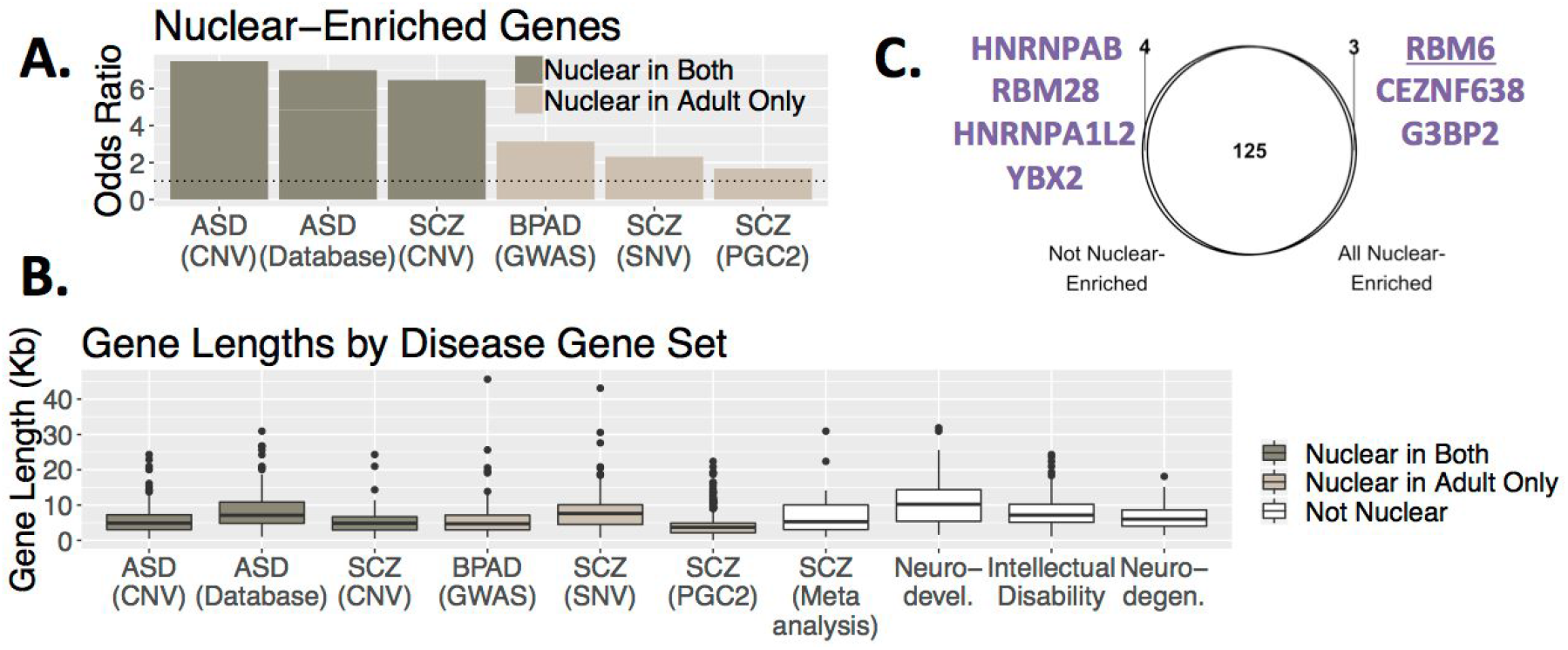
Disease-association. **(A)** Odds ratios for enrichment of genes significantly greater expressed in nuclear RNA in gene sets associated with neuropsychiatric diseases using Fisher’s exact test (FDR≤0.05). **(B)** Length in kilobases (kb) of genes in 10 gene sets associated with autism spectrum disorder (ASD), schizophrenia (SCZ), bipolar affective disorder (BPAD), neurodevelopmental disease (Neuro-devel.), intellectual disability, and neurodegenerative disease (Neuro-degen.). Where applicable, the type of study from which the gene association with disease derives is listed in parentheses. Colors indicate whether the gene set is significantly enriched for genes higher expressed in nuclear RNA in both ages or in adult cortex only (FDR≤0.05). *TITIN*, a near 118 kilobase (kb) gene in the ASD (Database) set that was 73.3 kb longer than the second longest disease-associated gene, is not depicted. (C) Overlap of RBPs with sequence motifs found (FDR≤0.01) in gene sets enriched in the nuclear compartment (listed in Fig 6A) compared to the four gene sets that were not. RBM6 was also exclusively identified in the nuclear-expressed gene set (see Figure 5A).

Because genes with neuronal functions as a group are longer than average (43) and longer genes are more abundant in the nuclear compartment, we checked if the genes in the disease sets that were over-represented in the nuclear-enriched genes were longer than other genes. While genes in the six nuclear-enriched sets listed in Fig. 6A were significantly longer than all expressed genes (t=25.7, FDR=2.9e-119), they were as a group significantly shorter than the genes in the other unassociated disease sets that were tested (t=-5.6, FDR=5.4e-08; Fig. 6B). Therefore although these genes were overall longer than average gene length, length cannot explain the association with nuclear localization.

We further explored the relationship between disease-associated gene sets and mechanisms that regulate RNA localization and found no difference in 3’UTR length or predicted 3’UTR minimum free energy between the nuclear-enriched disease-associated gene sets and those that were not (FDR>0.10; Fig S9A). There was also no difference in the number of A→I editing sites identified between the two groups of gene sets (FDR=0.14; Fig S9B). Interestingly however, introns within genes in the nuclear-enriched sets had greater IR ratios than in the other sets (t=7.4, FDR=5.0e-13; Fig S9C). In other words, disease gene sets enriched in the nuclear compartment had greater levels of intron retention. This result was not due to the nuclear-enriched sets having longer or more introns as disease gene sets that were nuclear-enriched in both ages or adult only had significantly shorter introns (t=-8.5, FDR=2.1e-17) and similar numbers of introns (FDR>0.49) than those in the other gene sets. Examining RBP motif enrichment in the disease associated genes showed that seven RBPs had motifs that were uniquely enriched in one group of gene sets vs. the other (FDR≤0.01; Fig. 6C). One of the three RBPs with a motif found in a nuclear gene set but not the non-nuclear sets—RBM6—was also specific to nuclear-enriched genes in adult cortex compared to cytoplasmic genes (Fig. 5A) and has been shown to localize to the nucleus and act as a splicing factor (44). Splicing (or lack thereof) may therefore be a mediating component to the increased localization of ASD-, SCZ- and BPAD-associated genes in the nucleus.

To explore these results in an expanded cellular context, we compared fraction profiles of cytoplasmic and nuclear RNA from cell lines sequenced by the ENCODE Consortium (5). Strikingly, H1, a human embryonic stem cell line, showed far fewer differences by fraction than the more differentiated cell types tested (9 of 51,502 genes differentially expressed at FDR≤0.05; Fig. S10A). SK-N-SH, a cell line derived from neuroblastoma, was among the most distinct by fraction (13,985 of 51,502 genes differentially expressed at FDR≤0.05). In the ENCODE data, genes with significantly greater nuclear expression when controlling for cell type (FDR≤0.05) were enriched for several disease gene sets, including ASD genes from CNV studies, BPAD genes, and SCZ genes from GWAS loci (OR>1.6, FDR<4.9e-2; Table S7). ASD genes from the SFARI database were enriched in genes from either subcellular compartment (OR>1.5, FDR<0.042).

Nuclear-enriched genes in individual cell lines, however, only variably held to this pattern: while the nuclear-enriched disease sets in our data (ASD and SCZ genes from CNV studies and SCZ and BPAD genes from GWAS) were also enriched in nuclear genes in the neuronal lineage SK-N-SH line (OR>1.7, FDR<3.9e-02), and neurodegenerative disease genes were enriched in SK-N-SH cytoplasmic genes (OR=2.5, FDR=3.3e-02), ASD, SCZ and BPAD genes were all enriched in cytoplasmic genes in endothelial HUVEC cells, for instance (see Table S7). Overall, many more nuclear-associated disease gene sets were enriched in nuclear genes in each cell line than in the non-associated sets or in cytoplasmic genes (Fig. S10B). These results indicate that subcellular localization of genes associated with psychiatric disease is prevalent not just in brain but also in many other cellular contexts, suggesting that regulation of nuclear export of these transcripts may be a feature across many cell types.

## Discussion

Here we have characterized a snapshot of RNA compartmentalization in developing and mature human postmortem prefrontal cortex. We find that despite the presence of pre-mRNA, the nuclear RNA compartment alone can be used as a relatively high fidelity stand-in for the whole transcriptome when focusing on gene-level expression. Both nuclear and cytoplasmic RNA captured similar numbers of differentially expressed genes between developmental stages, and the magnitude of change detected was highly correlated between fractions. The use of PolyA+ library preparation minimizes the difference between subcellular fractions; the proportion of splice junctions detected was comparable between fractions when measured using PolyA+ libraries, but significantly less so in nuclear RNA when measured with Ribo-Zero.

Interestingly, differences in expression between fractions were much more muted in prenatal compared to adult cortex. We identified over 47 times more genes differentially expressed by fraction in adult than prenatal cortex in PolyA+ samples. Transcription has been shown previously to be more widespread in prenatal brain than at more mature time points, with 4.1% of the prenatal genome transcribed compared to 3.1% of the adult genome (3). We also show that prenatal samples had a higher proportion of splice junctions, indicating that a greater volume of prenatal transcription is being processed. Given that the cellular composition of prenatal cortex includes a higher proportion of neural progenitor cells and embryonic stem cells and that these immature cells have a more plastic epigenome (45), it is tempting to speculate that as the brain matures, relative nuclear retention of RNA becomes a more utilized regulatory strategy in cells of the brain. This hypothesis is supported by the H1 embryonic stem cell line showing fewer differentially expressed genes by fraction than the other more differentiated cell lines profiled by ENCODE. It has also been shown that nuclear pore composition changes as cells differentiate and mature (19), so it may be that nuclear pores and transport mechanisms are less mature in fetal brain and passage from nucleus to cytoplasm is less restricted.

At the gene level, trends in developmental and subcellular compartment expression patterns suggest nuclear sequestration of developmentally down-regulated RNA. For instance, nuclear-enriched genes in the adult brain were relatively greater expressed in prenatal than adult in cytoplasm, but greater in adult than prenatal in nuclear RNA. Genes with changing age expression in only one fraction also showed an inverted preference for fraction localization: genes decreasing in expression with age in the cytoplasm were greater in adult nuclear RNA, while genes decreasing in the nuclear fraction with age were greater in adult cytoplasm. In other words, genes with greater cytoplasmic expression in one age tended to be higher expressed in that age, while nuclear-enriched genes tended to be higher expressed in the opposite age, suggesting that these nuclear genes, most of which are protein-coding, are being down-regulated at least in part by RNA not being exported to the cytoplasm for translation. While this pattern must be tested in single cell types to be confirmed, it suggests an added layer of regulation to be considered in the design of next-generation sequencing studies.

By profiling RNA editing across fractions and ages, we found that RNA editing was not globally associated with RNA localization by fraction, although we identified many sites that were unique to a fraction in one age that were found in every sample of that fraction and age group. These unique editing site groups were found in genes that were more highly edited than other genes and that were more highly expressed in those samples than in the opposite age or fraction, although almost all of the exons and introns targeted for editing were present in the other fraction or age in question in an unedited form. The limited read depth in the samples, however, challenges the conclusiveness of the RNA editing analysis. Future work that probes the relationship of these unique sites to localization and expression should study specific cell types at greater coverage.

IR has been shown recently to be a common splice variant type that increases during development in several cell types including neurons (12, 15, 16). Here we characterize splicing patterns across fractions in prenatal and adult cortex and confirm that although the majority of introns are constitutively spliced, IR is an abundant splice variant type—particularly in nuclear RNA—and increasing nuclear IR in a subset of introns correlates with increasing nuclear expression compared to cytoplasm and decreasing expression overall. Like global gene expression, specific splice variants seemed to pass more readily through the nuclear membrane in prenatal cortex than in adult, leading the decreasing relationship to be lacking in prenatal brain. It is unclear why increasing retention in age-associated introns is negatively correlated with expression only in prenatal cytoplasm samples, particularly as 28 of the 31 developmentally-retained introns were more retained in prenatal than adult samples in either compartment, counter to the expectation that increased IR could be sequestering these transcripts in the nucleus as the brain matures. Further work in specific cell types or single cells with increased sequencing depth will be required to resolve these relationships. Nevertheless, given that differentially retained introns by fraction were also more likely to be differentially retained by age, IR provides a link between developmental and compartmental expression trajectories in the data.

Identifying motifs of several known splicing factors with neurodevelopmental functions—such as RBFOX1 and RBFOX2—to be exclusively enriched by RNA fraction further strengthens the link between splicing and subcellular localization. We also find that 3’UTRs of nuclear-enriched genes are predicted to be more thermodynamically stable than cytoplasmic genes, suggesting that there may be a conformational similarity between these RNAs that leads to their increased nuclear expression. However, motif enrichment alone is a weak predictor of ***in vivo*** RBP binding, as many RBPs have been shown to bind similar motifs, but show more diversity in their preference for other supplementary factors such as neighboring sequence, RNA secondary structure, and use of bipartite motifs (35). Although future work will be needed to validate the role of these RBPs in regulating nuclear transcript levels, this preliminary work further implicates splicing to be a critical step in developmentally-regulated mRNA transport in human cortex.

Finally, we found that nuclear-enriched genes were also preferentially enriched in gene sets associated with neurodevelopmental psychiatric diseases but not with other brain diseases. Previous work has identified the importance of proper nucleocytoplasmic transport in brain diseases, particularly neurodegenerative diseases such as fronto-temporal dementia and amyotrophic lateral sclerosis (24, 25). Genes associated with these and related diseases were associated with increased adult cytoplasmic expression, in line with their important roles in mature cortex. Surprisingly, however, we found that genes associated with neurodevelopmental psychiatric diseases like ASD, SCZ, and BPAD were more likely to have higher expression in the nucleus at both ages tested, but particularly in adult where more nuclear sequestration in general was found. This association was not related to neuronal genes being longer and therefore taking more time to leave the nucleus. Curiously, this preference for nuclear localization of developmental neuropsychiatric disorder gene sets extended to other immortalized cell types profiled by ENCODE. The nuclear-associated gene sets also showed increased intron retention compared to the others and shared enriched RBP motifs associated with splicing. These results suggest that these genes may be undergoing extra processing or regulation in the nucleus that may make them more vulnerable to dysregulation, potentially mediated by alternative splicing and RBP binding and particularly during early adult life. That these sets of genes appear to have easier exit from the nucleus during fetal life is consistent with other data that expression of these disease gene sets tends to be greater during fetal than postnatal life (3, 42).

Nonetheless, this study is limited by lack of single cell or cell type-specific insight into these patterns. By using bulk human postmortem brain tissue, we trade improved clinical validity for reduced resolution of nucleocytoplasmic expression patterns. As mentioned previously, prenatal and adult cortices are populated by different cell types in different proportions, each with different proliferation, potency, and connectivity patterns that may influence the import-export decisions across the nuclear membrane. Cortical dissections were, however, matched for cell type composition at the tissue level within an age, lessening the potential for confounding by composition differences. Despite having to average the signal across cells and cell types, the fact that we still see this association between nuclear-expressed genes and psychiatric disease genes suggests that further study of this relationship is warranted.

## Acknowledgements

The authors would like to thank Jeffrey D. Rothstein for his insightful comments.

## Author Contributions

- A.J.P.: Conceptualization, Formal Analysis, Visualization, Writing – Original Draft Preparation, Writing – Review & Editing.
- T.H.: Formal Analysis.
- R.T.: Investigation.
- E.E.B.: Data Curation, Writing – Review & Editing.
- A.R.: Data Curation, Writing – Review & Editing.
- J.S.: Supervision, Writing – Review & Editing.
- T.M.H: Data Curation, Resources, Writing – Review & Editing.
- J.E.K.: Data Curation, Resources, Writing – Review & Editing.
- A.E.J.: Conceptualization, Supervision, Writing – Review & Editing.
- D.R.W.: Conceptualization, Funding Acquisition, Supervision, Writing – Review & Editing.

## Data availability and materials

The 12 PolyA+ library sample data are publicly available via BioProject (Accession PRJNA245228). The 11 RiboZero sample data will be made publicly available upon publication. Code is available through GitHub at: https://github.com/LieberInstitute/BrainRNACompartments.

## Materials and Methods

### Post-Mortem Brain Samples

Three prenatal and three adult human postmortem brains were selected from the collection of the Lieber Institute for Brain Development for use in this study. Brains in this collection were acquired, dissected, and characterized as described previously (3, 46). Briefly, post-mortem human brain was obtained by autopsy primarily from the Offices of the Chief Medical Examiner of the District of Columbia and the Commonwealth of Virginia, Northern District after informed consent from legal next of kin (protocol 90-M-0142 approved by the NIMH/NIH Institutional Review Board). Brain tissue was stored and dissected at the Clinical Center, NIH, Bethesda, Maryland and at the Lieber Institute for Brain Development in Baltimore, Maryland, and brain material was donated and transferred to the Lieber Institute under an approved Material Transfer Agreement. Post-mortem fetal brain tissue samples were provided by the National Institute of Child Health and Human Development Brain and Tissue Bank for Developmental Disorders (http://www.BTBank.org/) under contracts NO1-HD-4-3368 and NO1-HD-4-3383. The Institutional Review Board of the University of Maryland at Baltimore and the State of Maryland approved the protocol, and the tissue was donated to the Lieber Institute for Brain Development under the terms of a material transfer agreement. Clinical characterization, diagnoses, toxicological analysis, and macro- and microscopic neuropathological examinations were performed on all samples using a standardized protocol approved by the Institutional Review Board of the University of Maryland at Baltimore and the State of Maryland. Subjects with evidence of macro- or microscopic neuropathology, drug use, alcohol abuse or psychiatric illness were excluded.

### Cytoplasmic and Nuclear RNA Purification and Sequencing

A diagram of the study design is included in Fig. S1. Homogenate gray matter from the dorsolateral prefrontal cortex (DLPFC) approximating BA46/9 in adults and the corresponding region of PFC in prenatal samples were used for RNA extraction. To purify cytoplasmic from nuclear RNA, we used the Norgen Biotek Corp. Cytoplasmic and Nuclear RNA Purification Kit (Cat # 21000, 37400) following the manufacturer’s protocol including the optional DNase I treatment. RNA-sequencing libraries were prepared from each RNA fraction using PolyA-selection (“PolyA+”; Illumina TruSeq Stranded Total RNA Library Prep Kit, Cat #RS-122-2201) and rRNA-depletion (“Ribo-Zero”; Illumina Ribo-Zero Gold Kit (Human/Mouse/Rat), Cat # MRZG126) protocols to enrich for mRNA species. The resulting 24 libraries were then sequenced on one lane of an Illumina HiSeq 2000; the Illumina Real Time Analysis (RTA) module performed image analysis and base calling, and ran the BCL converter (CASAVA v1.8.2), generating FASTQ files containing the sequencing reads. “Br5339C1_polyA” and “Br5340C1_polyA” fastq files were downsampled to 24 million total reads to make the read depth more comparable across samples by joining paired read files, randomly shuffling read order while maintaining read pairs, and limiting the new downsampled FASTQ file to the top 12 million read pairs in the file.

### Data Processing and Quality Control

Raw sequencing reads were mapped to the hg19/GRCh37 human reference genome with splice-aware aligner HISAT2 (47) version 2.0.4, with an average 86.8% alignment rate for PolyA+ samples and average 92.6% alignment for Ribo-Zero samples. Feature-level quantification based on GENCODE (release 25, lift 37) annotation was run on aligned reads using featureCounts (subread version 1.5.0-p3) (48). Exon-exon junction counts were extracted from the BAM files using regtools (49) v. 0.1.0 and the ‘bed_to_juncs’ program from TopHat2 (50) to retain the number of supporting reads. Annotated transcripts were quantified with Salmon (51) version 0.7.2. Finally, alignment/processing metrics and the featureCounts results for genes, exons, exon-exon splice junctions, and annotated transcripts were read in and structured into analyzable matrices using R version 3.3.1. As a quality control check, raw FASTQ files were run through FastQC software (52), and all samples passed quality statistics including GC content, adapter content, and overall quality. After pre-processing, all samples passed additional QC checks for alignment rate, gene assignment rate, and mitochondrial mapping rates.

We downloaded ENCODE data in FASTQ format from the Gene Expression Omnibus (accession #GSE26284), and processed them using the same pipeline.

### Gene Expression Analysis

Principal component analysis was done using the plotPCA() function from the DESeq2 bioconductor package (53). Read distribution was calculated using the read_distribution.py function in the RSeQC suite (54). Annotation features were assigned in a prioritized order, so that reads overlapping coding (CDS) exons were labeled first, then untranslated (UTR) exons, then introns, and finally intergenic regions.

Gene expression differences were measured using the DESeq2 bioconductor package. Samples were segregated by library, fraction and age and compared using several linear models. Gene expression was first modeled by library type in the 11 nuclear samples using “~ Age + Library”. To check localization patterns of known nuclear and cytoplasmic genes, we modeled “~ Age + Fraction” separately in the 12 PolyA+ and 11 Ribo-Zero samples. Adult and prenatal samples from each library separately were assessed for differential gene expression by fraction (“~ Fraction”), while nuclear and cytoplasmic samples from each library were separately assessed by age (“~ Age”), culminating in eight sets of results.

For subsequent gene expression analyses, a gene was considered significantly differentially expressed if the absolute value of the log_2_ fold change in expression was greater than or equal to one, and if the false discovery rate (FDR) was less than or equal to 5%. We subset these genes according to whether they were in agreement across ages if measuring changes in expression by fraction, or across fractions if measuring changes by age, resulting in eight groups (e.g., both nuclear, both cytoplasmic, nuclear in prenatal only, nuclear in adult only, cytoplasmic in prenatal only, cytoplasmic in adult only, nuclear in prenatal but cytoplasmic in adult, and cytoplasmic in prenatal but nuclear in adult for comparison of gene expression by fraction). “Interaction” genes were considered those meeting the above criteria using the model “~ Age + Fraction + Age: Fraction” in the 12 PolyA+ samples.

Gene and disease ontology enrichments were calculated using the compareCluster() function from the clusterProfiler bioconductor package (55). To assess enrichment in neuropsychiatric disease gene sets, we used brain disease gene sets from Birnbaum *et al*. (42) and calculated enrichment of these genes within the nine groups of fraction-associated genes described in the previous paragraph (FDR≤0.05), after filtering disease gene sets for genes whose gene symbol were represented in the genes expressed in the dataset, using Fisher’s exact test.

ENCODE samples were first analyzed using the linear model “~ Cell type + Fraction + Cell type: Fraction” data in FASTQ format from the Gene Expression Omnibus (accession #GSE26284), and individual cell types were analyzed for fraction expression changes using the model “~ Fraction.”

Cell type specific expression patterns were detected using publicly available cell type-specific single cell RNA-seq data (33). The 466 single cell RNA-seq samples were processed as described above. Cytoplasmic genes included all genes within the “both cytoplasmic”, “cytoplasmic in prenatal only”, and “cytoplasmic in adult only” gene sets, and nuclear genes included all genes in the “both nuclear”, “nuclear in prenatal only”, and “nuclear in adult only” sets. For each gene, the cell type with the maximum reads per million mapped (RPM) was assigned as the cell type expressing that gene, and the proportion was calculated based on the total number of cytoplasmic or nuclear genes assessed.

Nucleoporin genes were identified via the AmiGO 2 Gene Ontology Database (http://amigo.geneontology.org/grebe) by searching Gene Ontology terms associated with the nuclear pore (GO:0005643, GO:0044613, GO:0044611, GO:0044614, GO:0044615, GO:0031080, GO:0070762, GO:1990876).

### Splicing Analysis

The proportion of splice junctions per sample were calculated by dividing the number of reads overlapping a known or predicted splice junction by the total number of reads. To characterize splice variant type use across the PolyA+ samples, we used the SGSeq bioconductor package (56). We first extracted features from the bam files using getBamInfo(), then used analyzeFeatures() to predict and quantify splicing events in each bam based on GENCODE (release 25, lift 37). We finally analyzed and summarized that output using analyzeVariants(), setting the minimum denominator to 10. The number of unique splice variants of each type were counted by extracting the types using variantType(). We calculated differential splice variant use by fraction and age using the DEXSeq bioconductor package (57). In building the DEXSeq dataset, we used the variant IDs as the featureID and the event IDs as the groupID in the DEXSeqDataSet() function. Similarly to the gene-level expression analyses, we subset the 12 PolyA+ samples by fraction and age and compared differential splice variant expression by fraction using the full model “~ sample + exon + fraction:exon” and the reduced model “~ sample + exon.” We compared splice variant expression by age using the full model “~ sample + exon + age:exon” and the reduced model “~ sample + exon.” We then stratified these results by splice variant type and used Fisher exact test to calculate the enrichment of each type in each fraction and age.

To further assess intron retention in the PolyA+ samples, we filtered introns from the IRFinder-IR-nondir.txt output of IRFinder (58) run on the Human-hg19-release75 reference for each sample. We excluded introns with the “NonUniformIntronCover” warning and those that had anything but “clean” listed in the GeneIntronDetails output column (i.e., excluding “anti-near”, “anti-over”, “known-exon+anti-near”, “known-exon”, and “known-exon+anti-near+anti-over”). Introns were further filtered to exclude introns with fewer than four reads spanning the splice junction or a junction using either the 5’ or 3’ exon-intron boundary, or with fewer than four reads supporting intron inclusion at each exon-intron boundary. To assess the relationship between gene expression and IR, we assigned the maximum IR ratio per sample for each gene from this filtered set of introns and compared IR ratios of genes regulated by fraction and age (FDR≤0.05) using Student’s t-test and Fisher exact test.

To quantify differential retention of individual introns, we subset the samples by fraction and age and filtered the IRFinder-IR-nondir.txt output to create four new lists, first filtering to only include the “clean” introns (from the GeneIntronDetails output column), then filtering constitutively spliced introns by group (i.e., adult, prenatal, nuclear, and cytoplasmic). We then used these new files as input to the analysisWithLowReplicates.pl function from IRFinder to calculate differential intron retention between fraction in prenatal and adult, and by age in nucleus and cytoplasm, using the Audic and Claverie test. We calculated the false discovery rate using p.adjust() and setting the n parameter to the total number of clean, non-constitutively splice introns in each comparison. The relationship between intron retention by fraction and age and gene expression was further examined by comparing counts of each using Fisher exact test.

Intron conservation was tested by extracting per base GERP scores for all “clean” introns from the UCSC Table Browser (hg19), calculating the mean score per intron, and comparing the means of groups of introns using Student’s t-test. Repetitive elements in introns were analyzed by downloading the RepeatMasker track from the UCSC Table Browser (hg19) and finding overlaps using findOverlaps() from the GenomicRanges bioconductor package (59).

### RNA Editing Analysis

RNA editing sites were called in the 12 PolyA+ samples as described previously (17). We annotated the RNA editing sites to genomic features using the GenomicFeatures bioconductor package (59) and a transcription database object built on GENCODE (release 25, lift 37). Overlap with repetitive sequences was assessed using the RepeatMasker track downloaded from the UCSC Table Browser (hg19) by finding overlaps using findOverlaps() from the GenomicRanges bioconductor package (59). We compared the editing sites identified in this study with previously identified editing sites using findOverlaps(). We examined the effect of fraction, age, and the interaction of the two on editing rate in the 1,025 sites present in all samples by first filtering the sites to those with a finite and non-NA logit-transformed editing rate in at least 5 samples and with at least one adult, prenatal, nucleus and cytoplasm represented and then using the model “~ Age + Fraction + Age:Fraction.” We compared the pattern of editing in our dataset of the 576 developmentally increasing editing sites from Hwang *et al*. (17) present using Fisher exact test.

We defined the sets of fraction- and age-specific editing sites by sites present in all samples of the listed first group that were not found in the second group. For instance, the “Adult Only” sites were present in all six adult samples but no prenatal samples. We assigned each editing site to the nearest gene using distanceToNearest() from the GenomicRanges package and compared the location of the site by fraction or age with the expression enrichment using the Fisher exact test. We identified KEGG pathway enrichment using compareCluster() for the ten groups of unique editing sites, setting the function to “enrichKEGG.” Annotation enrichment for these unique sites was also assessed using the Fisher exact test. To identify the major 3’UTR isoform, we identified which 3’UTR had the highest read coverage per gene.

### RNA binding protein motif enrichment analysis

We downloaded position weight matrices for human RNA binding proteins from the ATtRACT database (https://attract.cnic.es/; version 0.99B). Using makeBackground() function from the PWMEnrich bioconductor package (version 4.18.0), we calculated a lognormal background based on 1000 randomly selected gene cDNAs. For the RBP enrichment by fraction, we created FASTA files for four groups: cDNA for genes significantly higher expressed in nucleus or cytoplasm in adult or prenatal samples (FDR≤0.05). For the disease gene set analysis, we created FASTA files using cDNA for six groups: genes in sets nuclear enriched in both ages vs. those that were not, genes in sets nuclear enriched in adult only vs. those that were not, and genes in sets nuclear enriched in both OR in adult only vs. the four remaining gene sets. For each of the above 10 groups we called motifEnrichment() to calculate the enrichment for each motif, and then groupReport() to summarize the results over each gene set. RBPs with a motif that passed a significance threshold of FDR≤0.01 in the group report were considered enriched in the gene set.

### 3’UTR secondary structure and length analysis

We selected the highest expressed 3’UTR for each gene by annotating exons using the threeUTRsByTranscript() function from the GenomicFeatures bioconductor package (version 1.34.1) (59) and choosing the one with the highest mean expression across all samples per gene. We extracted cDNA sequence for these regions and calculated the minimum free energy using the RNAfold function in the ViennaRNA package (version 2.4.11). We assessed the difference between MFE in groups defined in the RBP motif methods section using the Student t test.

---

**Figure S1:**
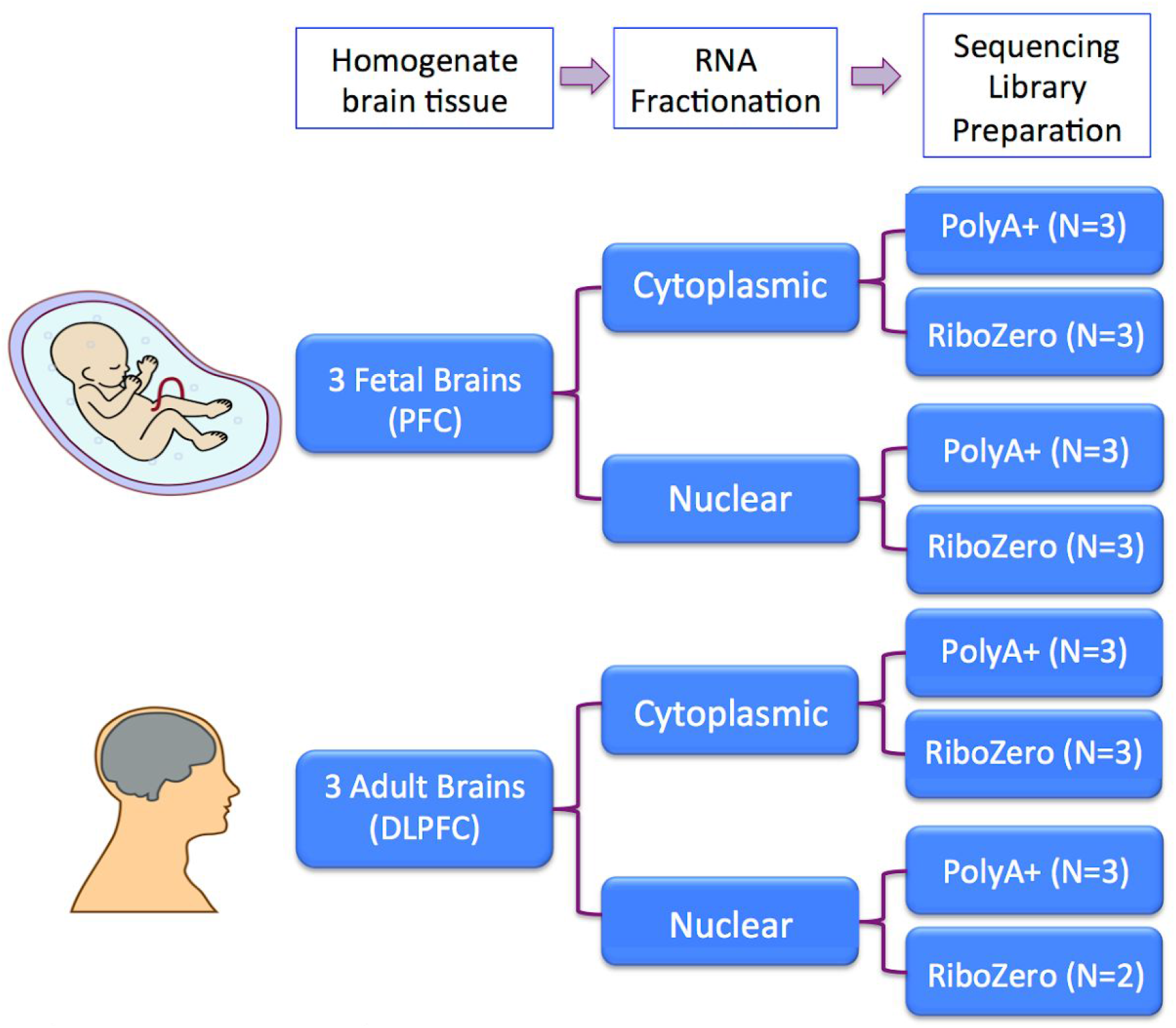
Experimental Design. We characterized the nuclear and cytoplasmic transcriptome in human prenatal postmortem prefrontal cortex (PFC) and adult postmortem dorsolateral prefrontal cortex (DLPFC) using two RNA sequencing library preparation methods. PolyA+ library preparation selects polyadenylated transcripts via a pull-down step, while RiboZero library preparation relies on a rRNA depletion step.

**Figure S2:**
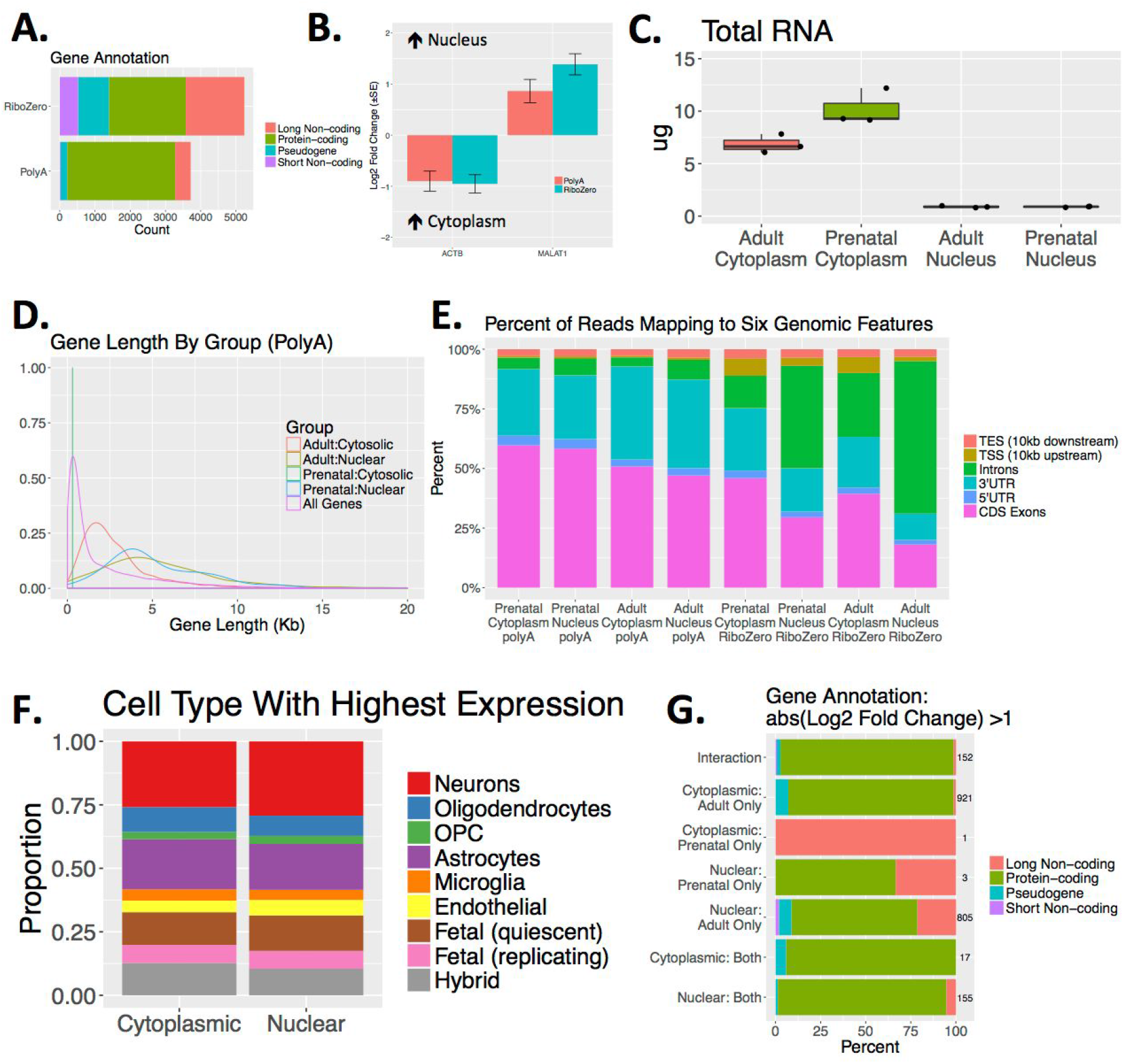
Characterizing the nuclear and cytoplasmic transcriptome in human brain. **(A)** Differentially expressed genes in nuclear RNA by library type (FDR≤0.05; abs(Log_2_ Fold Change)≥1). **(B)** *ACTB1*, a cytoplasmic gene, and *MALAT1*, a nuclear gene, are enriched in the appropriate subcellular fractions. Error bars reflect standard error. **(C)** Micrograms of RNA collected from each sample stratified by age and fraction. **(D)** Distribution of lengths of genes enriched by fraction in PolyA+ samples (FDR≤0.05; abs(Log_2_ fold change)≥1). “Cytoplasmic” and “Nuclear” reflect the fraction in which the gene is higher expressed, and “Adult” and “Prenatal” represent the age in which the comparison between fractions was made. **(E)** Percent of reads mapping to six genomic features in each group. TES=Transcription end site; TSS=transcription start site; UTR=untranslated region, CDS=coding; kb=kilobase. **(F)** The proportion of cytoplasmic- and nuclear-enriched genes (FDR≤0.05) maximally expressed in each cell type as measured in single cell RNA-seq data (33) **(G)** Annotation of groups of genes differentially expressed by fraction (FDR≤0.05; abs(Log_2_ fold change)≥1). The total number of genes in each group is listed to the right of each bar.

**Figure S3:**
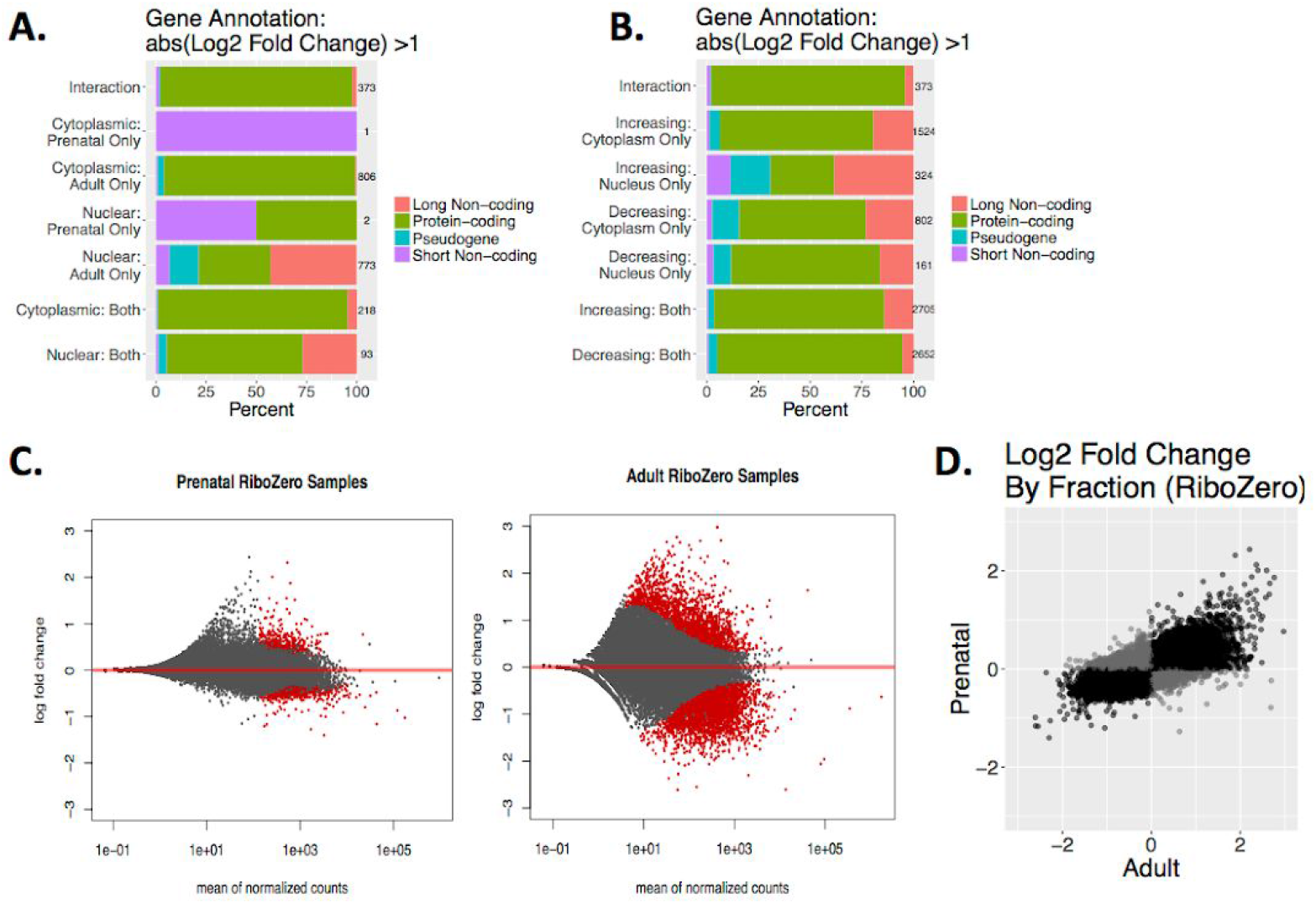
Comparing Fraction and Age in RiboZero Samples. **(A)** Annotation of groups of genes differentially expressed by fraction in adult and prenatal RNA (FDR≤0.05; abs(Log_2_ fold change)≥1). The total number in each group is listed to the right of each bar. **(B)** Annotation of groups of genes differentially expressed by age in cytoplasmic and nuclear RNA (FDR≤0.05; abs(Log_2_ fold change)≥1). The total number in each group is listed to the right of each bar. **(C)** MA plots of prenatal and adult gene expression differences measured across fraction. Red dots indicate FDR≤0.05. **(D)** Log_2_ fold change of expression across fraction in adult samples plotted against prenatal samples. Black dots indicate genes with agreeing sign, and gray indicate a change in Log_2_ fold change direction.

**Figure S4:**
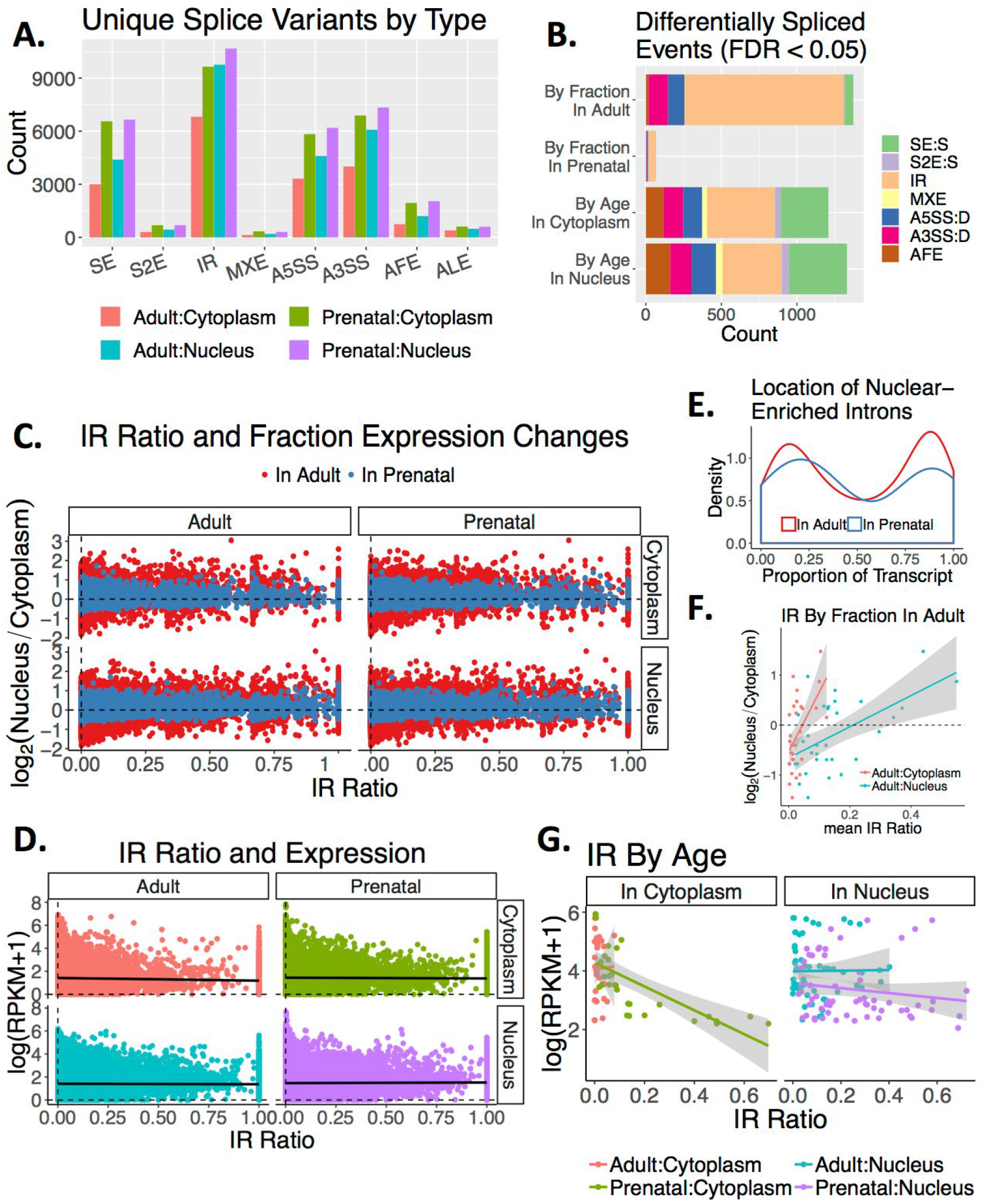
Alternative splicing patterns and intron retention. **(A)** Counts of unique skipped exons (SE), skipping of two exons (S2E), intron retention (IR), mutually exclusive exons (MXE), alternative 5’ exon splice site use (A5SS), alternative 3’ exon splice site use (A3SS), alternative first exon use (AFE) and alternative last exon use (ALE) stratified by fraction and age. Skipped exons (SE) and intron retention (IR) represented the greatest percentage of unique splice variants identified (50.8%). 72.9% more unique splice variants were identified in prenatal than adult samples, suggesting that the higher proportion of spliced RNA products in prenatal samples shown in Fig. 3A is accompanied by greater splicing diversity. By fraction, 42.8% more unique splice variants were identified in nuclear than cytoplasmic RNA. **(B)** Number of differentially expressed splicing events in different comparisons, stratified by splice variant type. As in overall gene expression, prenatal fractions showed fewer significantly differentially expressed splice variants than between adult fractions. Whether a splice variant was more expressed in nuclear than cytoplasmic RNA related to its variant type: significantly differentially expressed IR events by fraction (FDR≤0.05) were more likely to be more abundant in the nucleus (OR=50.9, FDR=8.7e-96), while SE, distal alternative 5’ exon start site (A5SS.D) and 3’ exon start site (A3SS.D) events were less abundant in the nucleus (OR=0.091, FDR<1.7e-07). **(C)** The log_2_ fold change of gene expression between fractions in adult (red) and prenatal (blue) plotted against the mean maximum IR ratio for an intron within each gene for each age:fraction group. Samples are stratified by panels indicating the age and fraction of the mean maximum IR ratio. 152,432 introns were shared between all 12 PolyA+ samples. Across samples, 58.68-85.33% of filtered introns were constitutively spliced in each sample, and 12.20-34.63% had an IR ratio (i.e., intronic reads divided by total intron and flanking exon reads) of greater than zero but less than five percent. **(D)** Gene expression measured as the log of the reads per kilobase per million mapped (RPKM) plus one plotted against the maximum IR ratio for each gene within each sample. Samples are colored and stratified into panels indicating their age:fraction group. The black line depicts the linear regression for each group. **(E)** Density plot of the location in a transcript of introns higher expressed in the nuclear compartment in adult and prenatal brain, listed as the proportion of the transcript length between the intron and the end of the transcript (3’<5’). Using the Audic and Claverie test for differential intron expression we found 35 introns significantly differentially retained within a transcript by fraction in adults, six by fraction in prenatal cortex, 10 by age in cytoplasmic RNA, and 21 by age in nuclear RNA (FDR≤0.05). Of the fraction-regulated introns, all were more retained (less spliced) in the nuclear compartment. These differentially retained introns tended to be single rather than clustered within a gene (90.4%) and were significantly shorter than the pool of total introns tested (t<-32.8, FDR<2.2e-25). Most relevant to this figure, fraction-regulated introns clustered at the beginning and end of the gene. **(F)** The log_2_ fold change of gene expression between fractions as measured in adult plotted against the mean IR ratio for introns differentially retained by fraction in adult (FDR≤0.05) in adult cytoplasmic samples (red) and adult nuclear samples (blue). The linear regression with standard error is shown for both groups. **(G)** The IR ratio of introns differentially retained by age when measured in cytoplasmic and nuclear samples, plotted by gene expression measured as the log of the RPKM plus one. Samples are colored by their age:fraction group. The linear regression line for each group is shown with shading to indicate the standard error.

**Figure S5:**
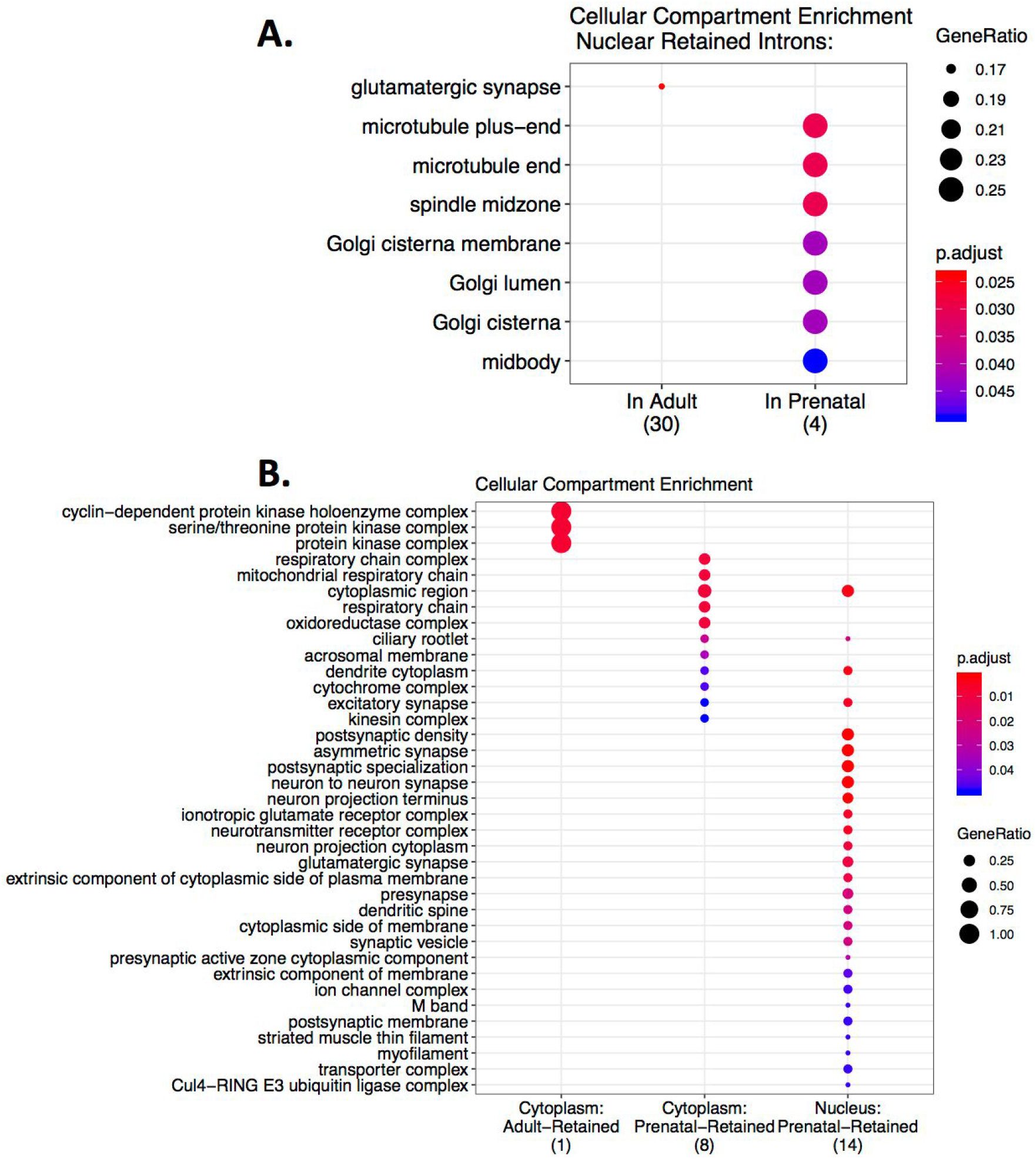
Cellular compartment ontology enrichment for retained introns. **(A)** Cellular compartment ontology terms enriched in genes containing introns that were higher expressed in nuclear than cytoplasmic RNA in adult (left) and prenatal (right) samples. **(B)** Terms enriched in genes containing introns more retained in adult than prenatal (far left) or more in prenatal than adult (middle and right columns) when measured in cytoplasm (left and middle columns) or nuclear RNA (right column).

**Figure S6:**
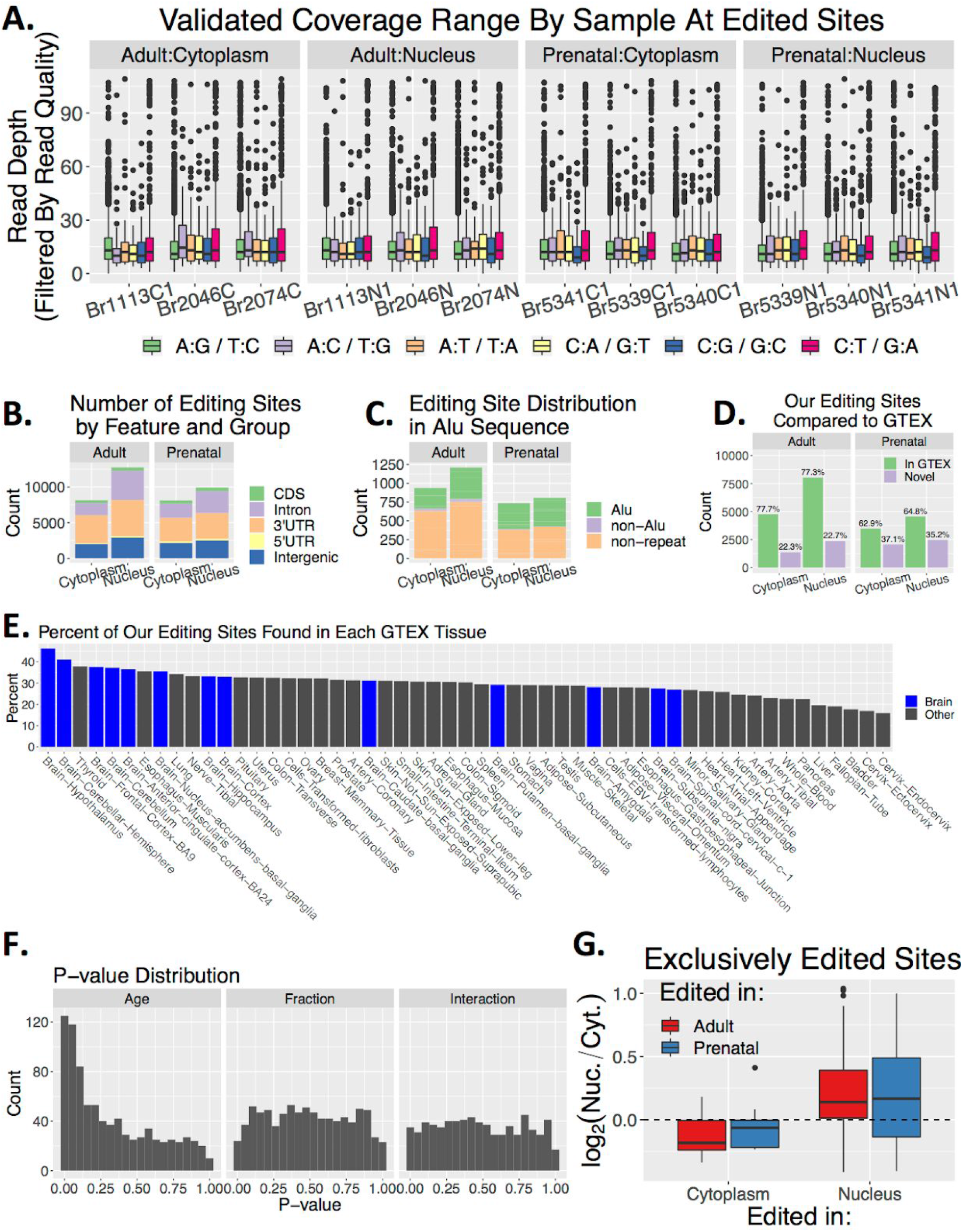
RNA editing across fraction and age. **(A)** Distribution of filtered read depth per sample for each editing context. “A:G/T:C” sites are considered A-to-I editing sites in our unstranded RNA-sequencing data. Read coverage was fairly even over all samples at edited sites, with a median coverage of 11-12 reads per site across samples. **(B)** Annotation of RNA features that contain an A-to-I editing site stratified by fraction and age. In line with previous reports, 21.7-33.8% fell within intronic sequence and 37.6-50.8% within 3’UTR sequence in each fraction and age tested. **(C)** Number of A-to-I editing sites that overlap an Alu sequence, a non-Alu repeat, or no repeats. 40.0-42.0% of A-to-I editing sites overlapped an Alu repeat sequence. **(D)** Number and percentage of A-to-I editing sites identified in our data that are also identified in GTEX consortium data. Of the previously unreported editing sites, 43.1% more were detected in nuclear than cytoplasmic RNA, and 13.8% more were detected in prenatal than adult samples. **(E)** Percentage of editing sites identified in each GTEX tissue that are also identified in our data. 69% of our 18,907 A-to-I editing sites were also detected in Genotype-Tissue Expression (GTEx) project data (60), particularly in GTEx brain samples (46.3%). **(F)** Distribution of unadjusted p-values calculated from linear regression assessing editing rate changes by age adjusting for fraction, fraction adjusting for age, or age:fraction interaction effects in the 1,025 sites found in all samples. After adjusting for false discovery rate, 81 sites were associated with age, while only 9 were associated with fraction and 6 with an interaction between age and fraction. **(G)** Log_2_ fold change of expression by fraction as measured in prenatal samples, in genes that include an editing site present in all cytoplasmic but no nuclear samples (right) or all nuclear and no cytoplasmic samples (left) in either adult (red) or prenatal (blue). The 159 adult nuclear editing sites were more likely to fall within intronic sequence than cytoplasmic sites (OR=5.9, FDR=9.8e-03), which were more likely to fall within 3’UTR sequence (OR=3.96, FDR=9.8e-3).

**Figure S7:**
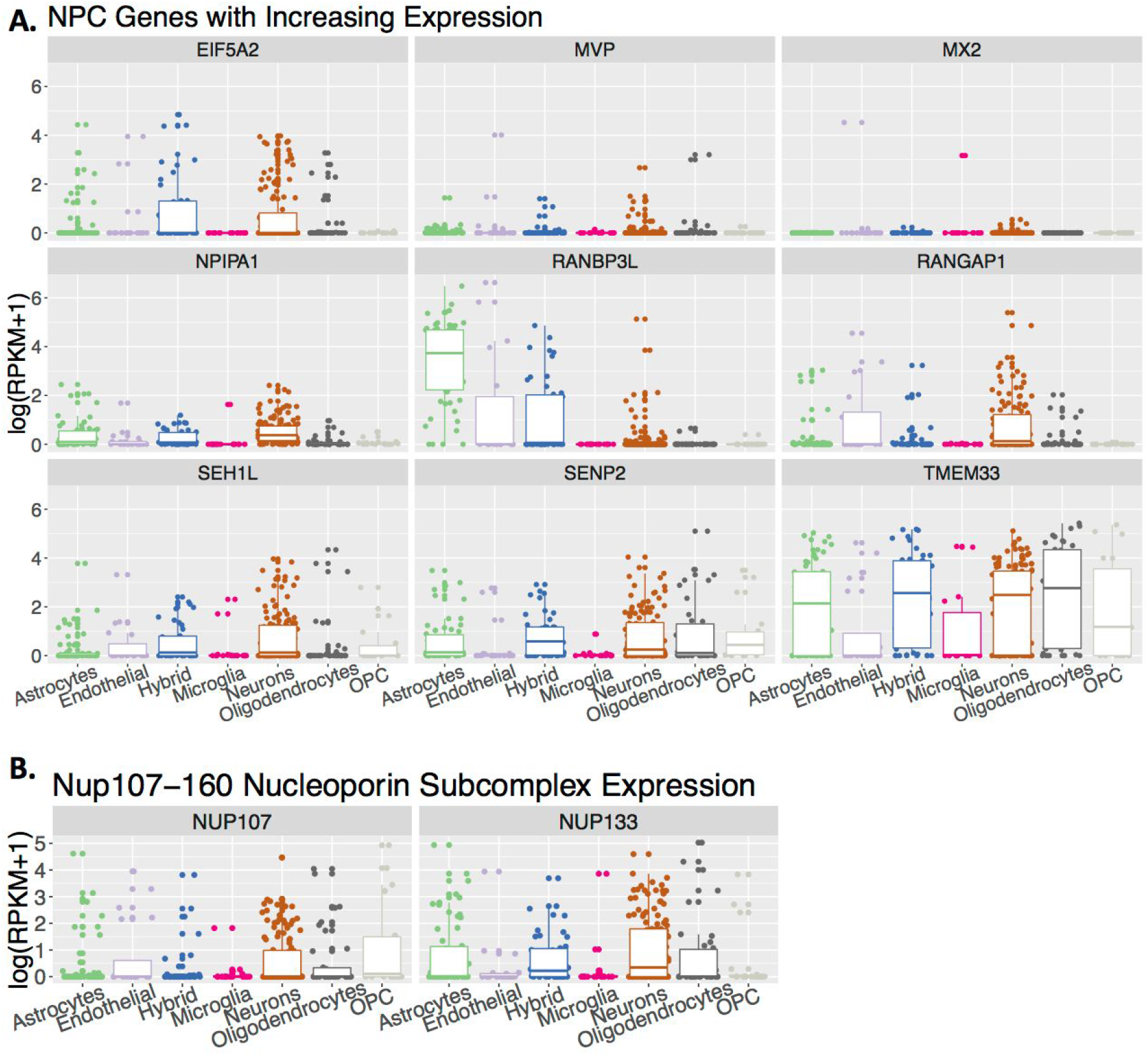
Cell type-specific expression of nuclear pore complex genes with increasing expression as the brain matures. **(A)** The log of the reads per kilobase per million mapped (RPKM) plus one of the nine genes with significantly increasing expression (FDR≤0.05) over cortical development as measured in a single cell RNA-seq dataset of cells isolated from adult human postmortem brain (33). **(B)** log(RPKM+1) of two components for the as measured in a single cell RNA-seq dataset of cells isolated from adult human postmortem brain (33).

**Figure S8:**
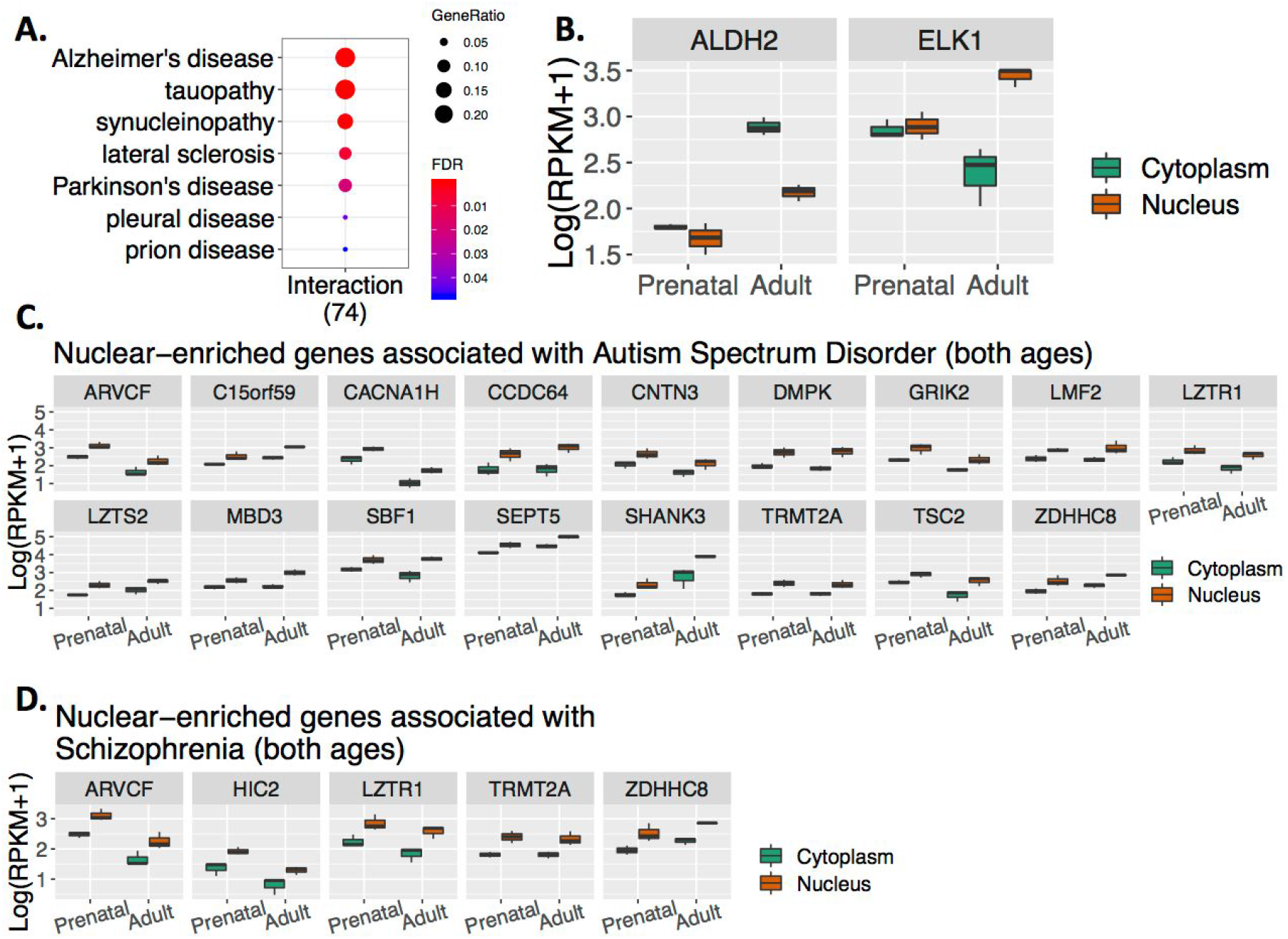
Disease Semantic and Ontology Enrichment. **(A)** Enrichment for disease ontology terms in “Interaction” genes whose expression varies by both fraction and age. Genes with a significant interaction between subcellular localization and age were enriched for involvement in Alzheimer’s disease and other neurodegenerative diseases (abs(log2 fold change)≥1; FDR≤0.05). **(B)** *ALDH2* and *ELK1* gene expression as measured in logarithm of reads per kilobase per million mapped reads plus one read (log(RPKM+1)), grouped by age (adult or prenatal) and fraction (cytoplasm and nucleus). Since the subcellular compartments are globally more similar in prenatal than adult samples, many of the “Interaction” genes were simply more highly expressed in adult than prenatal cortex overall, with greater expression in adult cytoplasm compared to nucleus and relatively similar expression between prenatal fractions. *ALDH2* is an example of greater expression in adult cytoplasm with muted prenatal compartmental differences. Some genes, however, such as the Alzheimer’s disease-associated gene *ELK1*, exhibited other patterns of interaction between fraction and age. Expression of *ELK1*—a transcription factor that regulates early action gene expression and is implicated in regulating chromatin remodeling, SRE-dependent transcription, and neuronal differentiation—was increased in adult nuclear RNA compared to the cytoplasm. In mice, Elk-1 protein abundance is tightly regulated by subcellular compartment as overexpression in the cytoplasm can lead to cell death (61). **(C)** Expression of autism-associated genes greater expressed in nuclear than cytoplasmic RNA in both adult and prenatal cortex (FDR≤0.05; abs(log_2_ fold change)≥1). **(D)** Expression of schizophrenia-associated genes that are greater expressed in nuclear than cytoplasmic RNA in both adult and prenatal cortex (FDR≤0.05; abs(log_2_ fold change)≥1).

**Figure S9:**
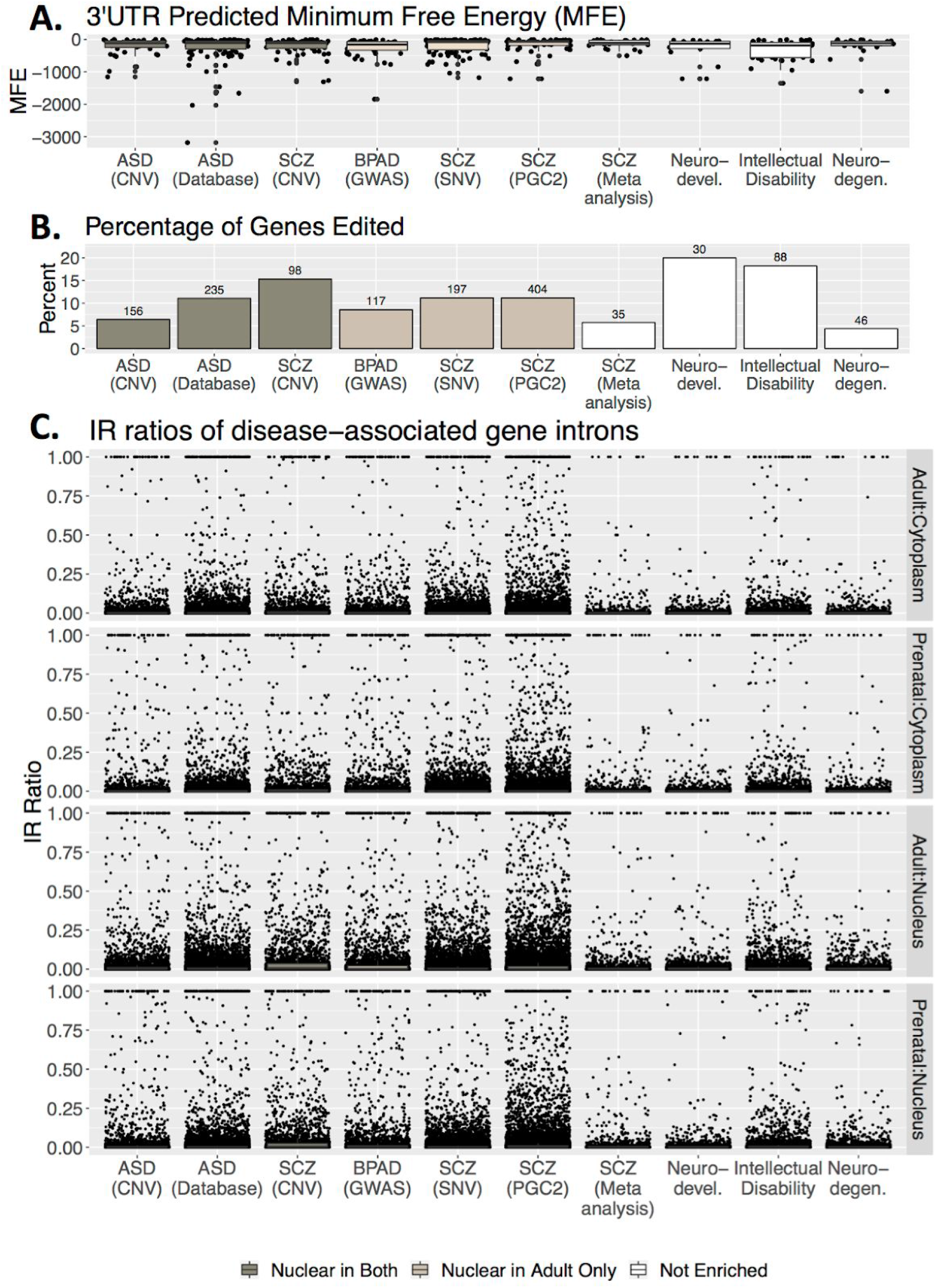
Mechanisms of subcellular localization and disease-associated genes. **(A)** Minimum predicted free energy of the highest expressed 3’UTR for each gene in each set associated with psychiatric diseases. **B)** The percentage of genes in each set that were found to have an A→I editing site. The total number of genes in each set is listed above each bar. **C)** Intron retention ratio for each intron within a gene associated with psychiatric disease. Gene sets are color-coded according to whether they are enriched for genes that are greater expressed in the nuclear compartment in both adults and prenatal cortex, in only adult cortex, or neither.

**Figure S10:**
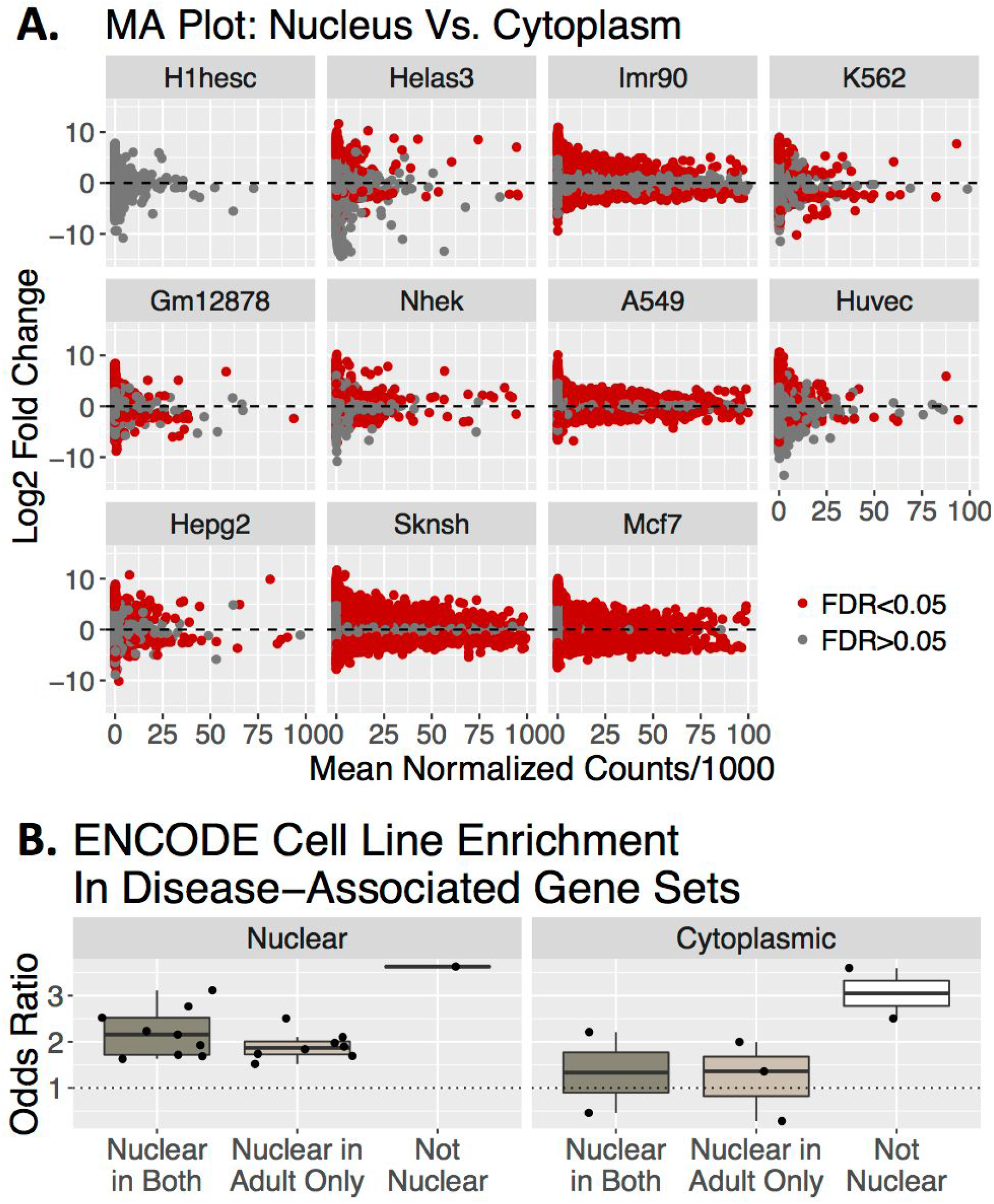
ENCODE Sample Disease Gene Set Association. **A)** MA plot showing the log_2_ fold change between fractions (positive values indicate greater expression in nucleus) and mean normalized counts divided by 1000 for the 11 cell lines profiled by ENCODE. Red dots indicate FDR≤0.05. **B)** Odds ratio for enrichment using Fisher’s exact test (FDR≤0.05) of nuclear- (left) or cytoplasmically-enriched genes (right) in the 11 ENCODE cell lines in the 10 disease-associated gene sets color-coded by whether the set is also enriched in the nuclear genes from our cortical data in both ages, adult only, or neither age.

